# Willingness to wait covaries with endogenous variation in cortisol

**DOI:** 10.1101/2021.09.18.460891

**Authors:** Evgeniya Lukinova, Jeffrey C. Erlich

**Affiliations:** NYU-ECNU Institute of Brain and Cognitive Science at NYU Shanghai, China; NYU Shanghai, Shanghai, China; N/LAB, University of Nottingham, Nottingham, UK; Shanghai Key Laboratory of Brain Functional Genomics (Ministry of Education), East China Normal University, Shanghai, China; Sainsbury Wellcome Center, University College London, London, UK

**Keywords:** cortisol, perceived stress, decision making

## Abstract

Stress is a normal part of our everyday lives. It alerts us to changes in our environment working as an early warning system. However, when stress is prolonged, it can become harmful. The deleterious effects of stress on brain function are well established: chronic stress significantly impairs cognitive function reducing our ability to solve problems and to regulate behavior and, therefore, may lead to more challenges that can further exacerbate stress. An important class of decisions that may be made under stress includes those between rewards delivered immediately vs. those delivered in the future. Not considering or devaluing future outcomes (delay discounting) can result in adverse outcomes such as not buying health insurance, gambling or drug use. To date, however, little is known about how chronic stress influences economic decisions that differ in the time of outcome delivery. A handful of studies suggest that increased stress may lead to more impulsive choices in subjects of average socioeconomic status and stress levels. Here, we addressed this gap by using a longitudinal design to determine whether chronic stress measures (via questionnaires, saliva and hair samples) are associated with the economic choices under different time scales within subject (N = 34). We found that the degree to which people think it is worth waiting, i.e. individual’s discount factor, over seconds, but not over days, varied reliably with endogenous stress. These results are imperative to studying stress in a variety of contexts: e.g., the variation in consumer’s impulse purchases could be attributed to the variation in individual stress responses to the environment, even in healthy adults.

## 1 Introduction

Prolonged stress is known to have negative influence on physical health (Harvey et al., 2003; Geronimus et al., 2010; Association et al., 2021) and psychological well-being (Faravelli and Pallanti, 1989; Hammen, 2005; Kendler et al., 1999; Kessing et al., 2003). Negative effects of stress on behavior can be seen through a range of maladaptive behaviors, such as poor emotional control and social withdrawal. There are a few dominant theories of how stress impacts the quality of decision making that distinguish between effects of short-term (or acute) and long-term (or chronic) stress (Kandasamy et al., 2014; Haushofer and Fehr, 2014; Mullainathan and Shafir, 2013; Cohn et al., 2015; Lerner et al., 2012; Riis-Vestergaard et al., 2018). Many studies find no significant effect of acute stress on economic choices (Sokol-Hessner et al., 2016; Lempert et al., 2012). Contrary, controlled state of chronic stress in a laboratory experiment significantly increased risk aversion and led to steeper delay discounting (Kandasamy et al., 2014; Riis-Vestergaard et al., 2018). Field data also support a correlation between chronic stress and poor economic decisions. For example, chronically-stressed individuals carry high-interest loans, have higher incidences of substance abuse, display a lower willingness to take risks and forgo current income for larger future incomes (Haushofer and Fehr, 2014; Mullainathan and Shafir, 2013). More so, stress may cause maladaptive and dangerous behaviors, making this study relevant to the prevention of substance dependence and various impulsivity disorders (Linsky and Straus, 1986; Lempert et al., 2012, 2018b; Lopez-Guzman et al., 2019).

We define stress as the “organism’s reaction to environmental demands exceeding its regulatory capacity” (Haushofer and Fehr, 2014). When a stressor is perceived, the hypothalamic-pituitary-adrenal-axis (HPA-axis) is activated and eventually releases cortisol (a steroid hormone, Schepers and Markus, 2015). Cortisol influences, regulates, or modulates many of the changes that occur in the body in response to stress. Hydrocortisone tablets work as a hormone replacement for a natural hormone called cortisol and are used in laboratory experiments to mimic the state of chronic stress (Kandasamy et al., 2014; Riis-Vestergaard et al., 2018).

Several common measures of chronic stress used in the recent literature include questionnaires (Holmes and Rahe, 1967; Goodman et al., 1998), collecting heart rate and blood pressure (Gooding et al., 2015), hair (Ceccato et al., 2015; Karlén et al., 2011), and saliva samples (Pruessner et al., 2003; Inder et al., 2012; Kandasamy et al., 2014). Stress questionnaires can track self-reported ‘stress’ traits or measure perceived stress. Saliva samples provide a measurement of the cortisol concentration at a single point in time (Hellhammer et al., 2009). Still, instead of tracking the peak of the diurnal cycle it is also possible to observe the level of cortisol over several days at the same exact time (Inder et al., 2012; Dietrich et al., 2013; Meggs et al., 2016; Pruessner et al., 2003; González-Cabrera et al., 2014) making it a better proxy for chronic stress (Lenow et al., 2017; Weckesser et al., 2019). Compared to all other biological stress measures hair samples have certain advantages, including its ability to be stored and transported at room temperature and to reflect cortisol levels over the period of hair growth (Karlén et al., 2011). Of course, all measures of stress are prone to be collected with some error. Having several measurements in this study and being able to extract a few, allows for better control and validation of the measurement system.

In this longitudinal study the main goal was to explore whether economic choices under various time scales could be accounted for by variation in stress over time (e.g., from negative income shocks or from non-economic stressors, like divorce, health problems, etc.) Following the existing literature, we hypothesized that we would find the positive relationship between behavioral impulsivity and bio-or self-reported stress: an individual with a higher stress level will consider less alternatives in the future (Riis-Vestergaard et al., 2018; Wang et al., 2020). We also hypothesized that individual fluctuations in stress levels would be associated with similar trends in decision making, i.e. a subject’s increase in stress level from one experimental session to the next would be accompanied by more impulsive choices in the later session (not confirmed by experimental studies so far, Lempert et al., 2018a).

Time preferences, which involve a trade-off between a larger outcome received later and a smaller outcome received sooner, have been found to predict various outcomes, including SAT scores, income, and anti-social behavior (Golsteyn et al., 2014; Åkerlund et al., 2016). In the laboratory, time preferences are typically assessed using the delay-discounting task. Our previous studies have demonstrated strong correlations and stability in time preferences for offers in seconds and days (Lukinova et al., 2019; Lukinova and Erlich, 2021). However, human delay discounting in seconds remains understudied, despite its potential for cross-species comparisons. The standard delay-discounting task involves decisions about outcomes occurring in days or weeks. In contrast, the delay-discounting task in seconds requires subjects to actively wait for the reward during each trial, rather than simply postponing the decision (building upon the distinction made by Paglieri, 2013). With our study’s main goal in mind we tested whether costs of waiting or postponing shared some intersubject or/and intrasubject variability with endogenous stress.

Here, we addressed the limitations of the previous literature and used a battery of incentivized decision-making tests and stress measures in order to find out whether chronic stress and its variation account for any variability in economic choice. To our knowledge, this is the first attempt to bring together in a within-subject longitudinal study using a sample of healthy working adults.

- a previously validated set of incentivized delay-discounting tasks across time scales;
- bio stress markers: hair cortisol and salivary cortisol; and
- self-reported stress measures

First, we explored time preferences and chronic stress measures separately. Second, we studied the similarity of distinct stress measures to delay discounting via correlation analysis. Third, we constructed predictive models by separating effects of stress on within-subject and between-subject behavioral variability. For the within-subject analysis, we used the changes in biological stress markers over time to predict the changes in delay discounting over time (in seconds and in days, separately). For the between-subject analysis, we related the biological stress markers and self-reported stress measures to subjects’ discount factors in seconds and in days. Finally, we checked whether we can accurately classify subjects of similar behavior profile by incorporating their stress measures. We acknowledge that our sample size is rather low for a (predominantly) correlational study. Nevertheless, we believe that using a unique set of stress measures together with behavioral measures allowed us to improve the scholarship on how time preferences interact with hormonal and neuromodulatory systems.

## 2 Results

Subjects were recruited from the general population of Shanghai, China (N = 41, 34 with all stress measures; Materials and Methods). They completed questionnaires, participated in a battery of decision-making tasks, and provided biological samples over four sessions scheduled every other week (Figure 1). The first three sessions that included behavioral tasks were scheduled at approximately the same time per each subject in order to ensure that salivary cortisol we collected can represent chronic stress. We collected one saliva sample per participant per session and one hair sample per participant two weeks after session 3 (analyzing three 1-centimeter segments separately to map onto stress over the last three months, Materials and Methods). At the beginning of the first session (but after providing the first saliva sample) subjects also completed stress questionnaires resulting in three self-reported stress measures.

**Figure 1.**
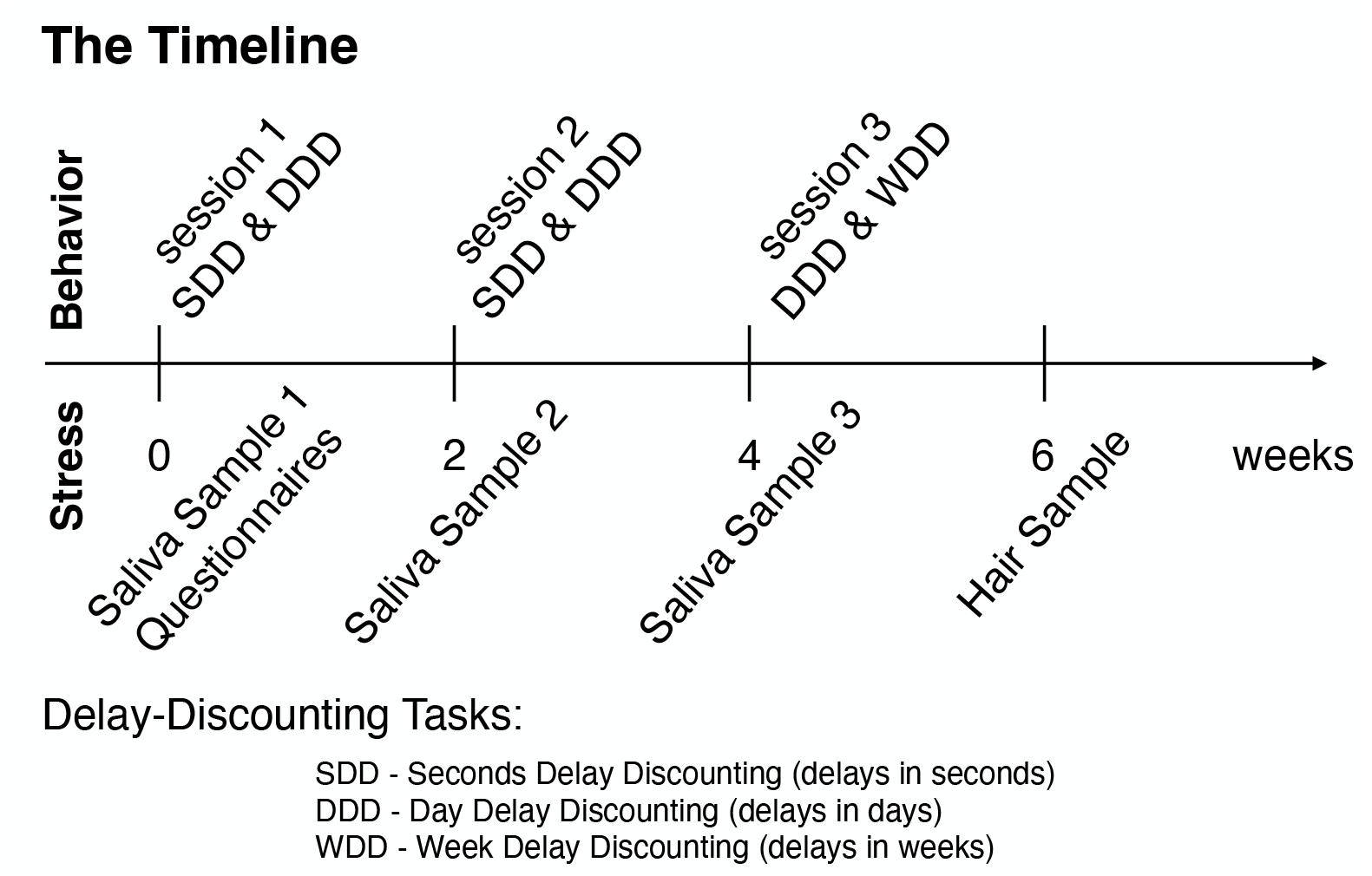
Experimental timeline. Each session took place every other week. During session 1 subjects provided the first saliva sample (by chewing on a synthetic swab), filled in the questionnaires, and participated in seconds and day delay-discounting (SDD and DDD, respectively, in counter-balanced order per subject) tasks on computers. During session 2 subjects provided the second saliva sample and participated in the same tasks as in session 1. During session 3 subjects provided the third saliva sample and participated in day and week delay-discounting (DDD and WDD, respectively) tasks. In two weeks after session 3 subjects returned to provide the hair sample (hair strands with the total thickness of a toothpick and at least 3cm in length cut twice).

### 2.1 Delay-discounting behavior is stable and reliable

In each session, subjects completed a series of delay-discounting tasks (Materials and Methods). During the first two sessions, subjects participated in tasks with delays in seconds and days (seconds delay discounting (SDD) & day delay discounting (DDD): 200 trials per task, Figure 1). During the third session, subjects participated in tasks with delays in days and weeks (DDD & week delay discounting (WDD): 100 trials per task, Figure 1). During each session, subjects experienced an alternating set of tasks: e.g., for the first two sessions (SDD-DDD-SDD-DDD, with the order counter-balanced per subject). In each trial of delay-discounting task, subjects made a decision between the sooner and the later options (Figure 2A). Only in the seconds delay-discounting task (with delays in seconds), subjects experienced the delay whenever delayed option was chosen at the end of each trial. Also, rewards from each trial were accumulated. In contrast, for other delay-discounting tasks (with delays in days or in weeks), subjects saw their choice confirmation and then proceeded to the next trial. For these tasks, at the end of the session, a single trial was selected randomly to determine the payment. To summarize, across three experimental sessions subjects participated in three distinct tasks: SDD, DDD, and WDD with delays in seconds, days, and weeks, respectively.

**Figure 2.**
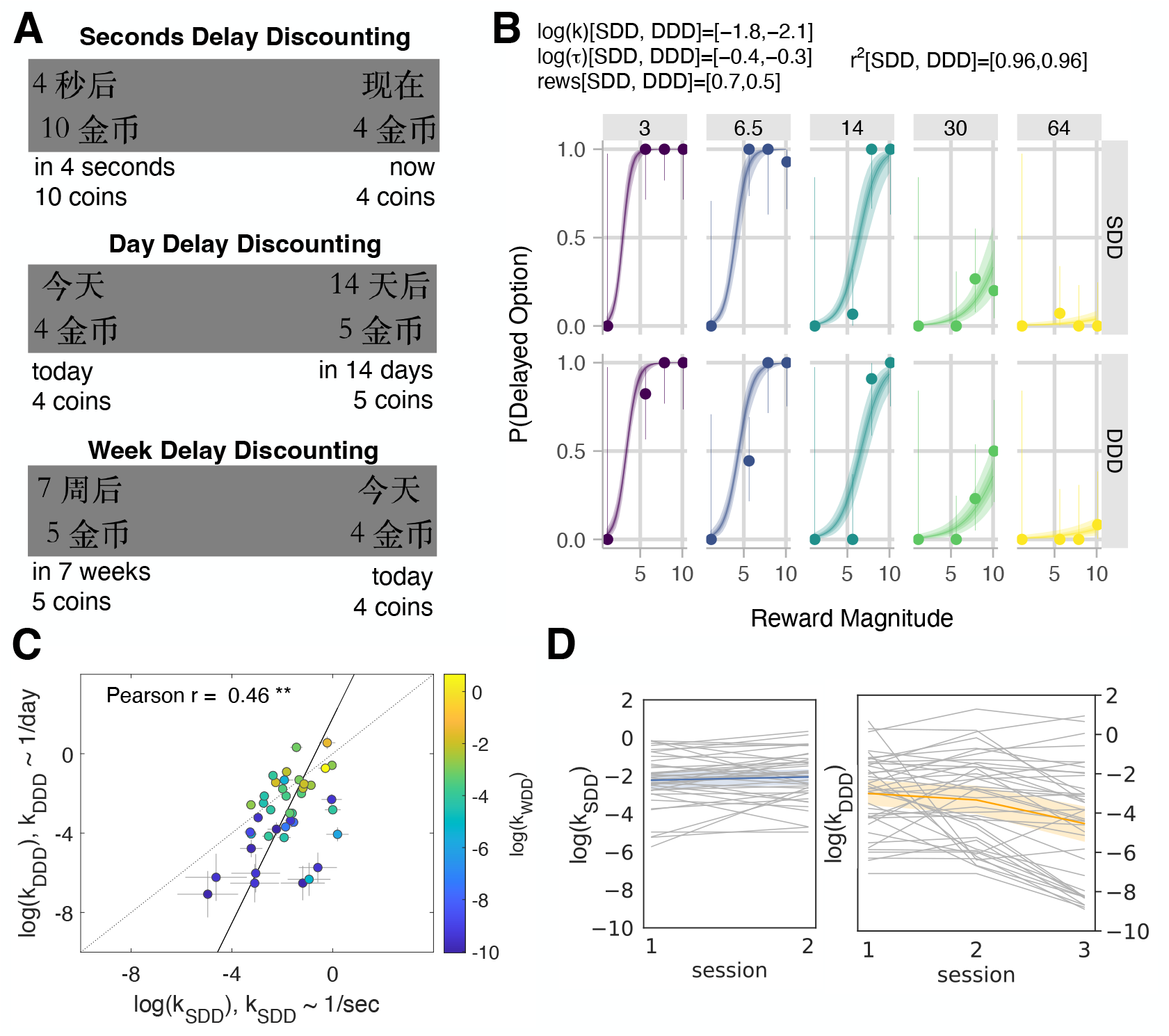
Comparison of model-based parameters across tasks: SDD (sessions 1 & 2 with delays in seconds), DDD (sessions 1, 2 & 3 with delays in days), and WDD (session 3 with delays in weeks). (A) Stimuli examples in the delay-discounting tasks through screenshots as presented to subjects in Chinese (Mandarin) and translation into English below. The ‘now/today’ option appeared on the left or the right part of the screen according to the random draw. (B) Model fit of an example subject (for SDD & DDD in sessions 1 & 2 combined).In each panel, the marker and error bar indicated the mean and binomial confidence intervals of the subject’s choices for that offer. The smooth ribbon indicated the BHM model fits (at 50, 80, 99% credible intervals). At the top of the subject plot we reported the mean estimates of model parameters for each task for that subject. We also listed the Bayesian *r*^2^ for each task. (C) Time preferences across time horizons. Each circle was one subject (*N* = 41). Discount factors were estimated in days but then converted to the units of the task. The error bars were the SD of the estimated coefficients. The solid line represented the perpendicular (or total) least squares (TLS) regression line (Huang et al., 2013). The dotted line was the unity line (*y* = *x*). The logs of discount factors (log(*k*)) in seconds delay task (SDD, sessions 1 & 2, x-axis) were plotted against the logs of discount factors in day delay task (DDD, sessions 1 & 2, y-axis). The color of the circles aligned with the logs of discount factors in week delay task (WDD, session 3) represented via a colorbar. Pearson correlation reported on the plot (*r* = 0.46, *p* = 0.003; ** for *p* < 0.01). (D) Stability of delay-discounting behavior across sessions in SDD (left) and DDD (right). Each subject’s log(*k*) (y-axis) was plotted via a line plot (with x-axis being an experimental session). The mean and 95% confidence interval were plotted using color.

Subjects’ time preferences were estimated by fitting their choices (24,277 choices across all experimental sessions) with a Bayesian hierarchical model (BHM) of hyperbolic discounting with decision noise and reward scaling per experimental session (Materials and Methods). In this paper, we followed previously validated methodology (Lukinova et al., 2019). The main model was compared to two additional models that reflected the reduced number of parameters. Based on 10-fold cross-validation criteria all comparisons favored the main ‘reward scale’ model (Materials and Methods and Table S1, SI). The main model resulted in subject fits comprised of mean estimates of log(*k*) -logarithm of the delay-discounting factor (discount factor), log(*τ*) - logarithm of the decision noise and reward scale parameters for each subject per experimental session and two delay-discounting tasks within it. Subject fits and correlations between fits across sessions and tasks were robust to the model choice (Table S2, SI). The subjects’ choices were well fit by the model as assessed using Bayesian *r*^2^ and reported in each of the subject delay plots (Figure 2B for an example subject fit; see SI for all subject plots in Figures S2 & S3 and posteriors in Figure S4). Consistent with previous scholarship (Lukinova et al., 2019; Lukinova and Erlich, 2021), we found that time preferences measured through discount factors showed strong correlations across different time horizons: between days and seconds (Pearson *r* = 0.46, *p* = 0.003, Figure 2C) and between days and weeks (Pearson *r* = 0.96, *p* < 0.001). Across sessions only log(*k*) in DDD was significantly different between session 3 and the first two sessions according to permutation tests (e.g., log(*k*): *M*_*s*1_ = − 3.002 & *M*_*s*3_ = − 4.534, *p* < 0.001, Materials and Methods). On the contrary, the economic behavior in both SDD and DDD tasks between session 1 and session 2 was stable (permutations tests for log(*k*) difference were not significant, *p* > 0.1, also in Figure 2D). Subjects who increased their patience over the gained experience with the task (i.e., for whom log(*k*) in session 3 was smaller than the average between sessions 1 & 2) were further explored.

### 2.2 Chronic stress variables convey distinct meanings

Prior to and after the delay-discounting tasks stress levels were measured using different methods (Figure 1, Materials and Methods). Importantly, we did not stress the subjects. Instead, we collected the base level of stress using i) Perceived Stress Scale (PSS, Cohen et al., 1983), The Brief Encounter Psychosocial Instrument (BEPSI, Frank and Zyzanski, 1988), and The Social Readjustment Rating Scale (SRRS, Holmes and Rahe, 1967) questionnaires before the decision-making tasks during session 1 (forming ‘pss’, ‘bepsi’, and ‘lcu’ variables, respectively, in Figure 3, Materials and Methods); ii) saliva samples before the decision-making tasks for each session scheduled at approximately the same time and the same day of the week per participant (three samples per three sessions: log-transformed cortisol levels ls1-ls3 in Figure 3); and iii) hair samples two weeks after session 3 (three 1cm-segments analyzed separately: log-transformed cortisol levels lh1, lh2 and lh3 from the stress level two months before collection to one-month-before and at the month of collection, respectively, see Figure 3). Due to the voluntary nature of sample collection, we only had 34 subjects with all stress data (of the 7 subjects with missing data, the majority were missing hair samples).

**Figure 3.**
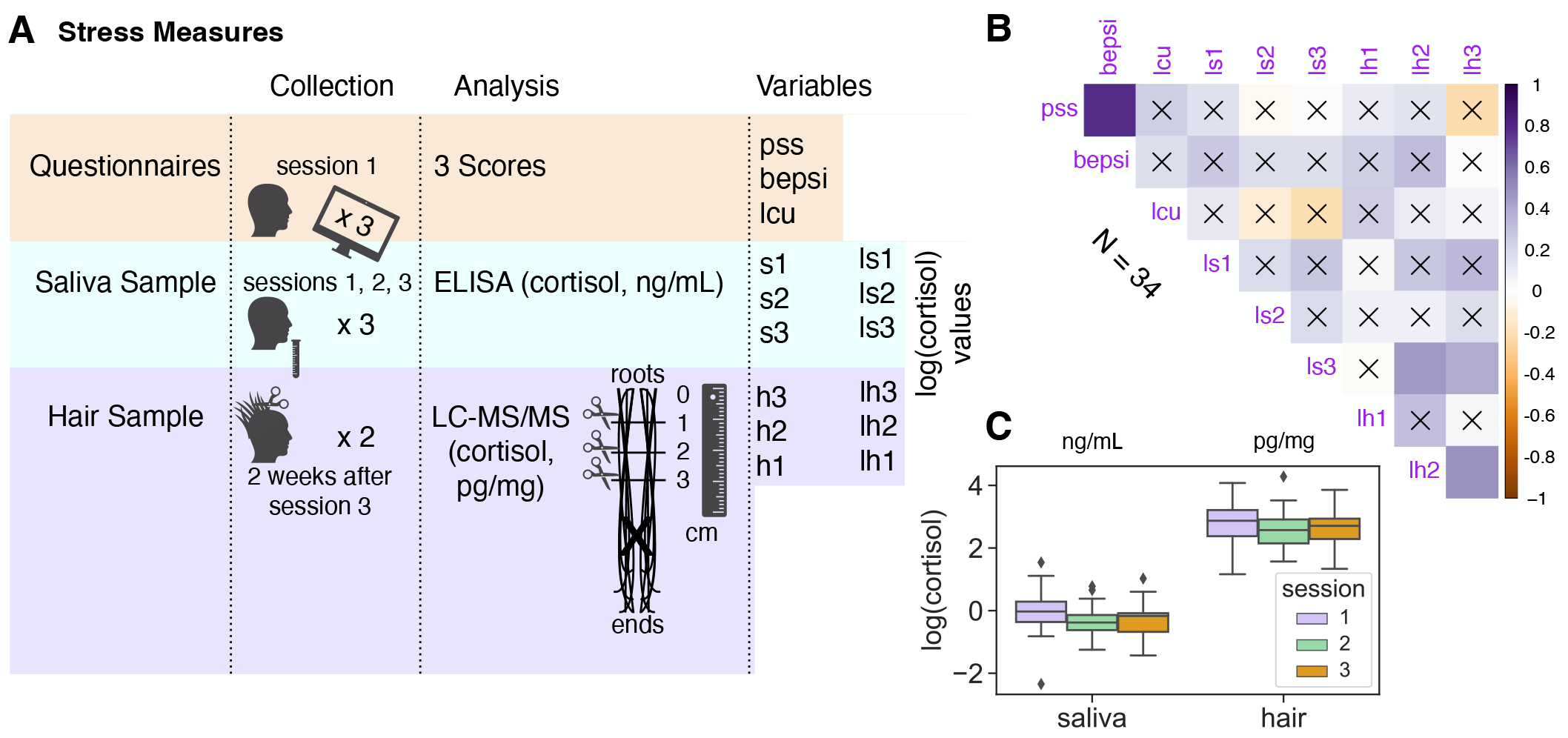
(A) Stress measures collection, analysis, and related variables (see details in Materials and Methods). Subjects filled in three questionnaires in session 1: Perceived Stress Scale (PSS, Cohen et al., 1983); the Brief Encounter Psychosocial Instrument (BEPSI, Frank and Zyzanski, 1988); and the Social Readjustment Rating Scale (SRRS, Holmes and Rahe, 1967) to get life change units. Also, subjects provided three saliva samples in total, one in the beginning of each experimental session. Finally, most of the subjects returned two weeks after session 3 to provide a hair sample (hair strands with toothpick thickness at least 3 cm in length cut twice were merged from the cut and then divided into three 1cm-segments reflecting the chronic stress two months, one month prior to and at the month of collection). (B) Correlation plot across all stress measures. The color intensity (vertical colorbar) signified the correlation *r* coefficient. The insignificant correlations, e.g. where *p* >= 0.05, were indicated with an ‘x’. (C) Boxplots of log(cortisol) for saliva and hair samples across three experimental sessions.

Stress questionnaires can track ‘stress’ traits, saliva samples may reflect level of cortisol over several weeks (when collected at the same time and on the same day of the week) and cortisol of hair samples divided into 1cm length segments can correspond to chronic stress (one month of stress per one 1cm segment following Gow et al., 2010). According to the literature, we expected significant medium level correlation between hair and saliva measures around *r* = 0.4 (Vanaelst et al., 2012; van Holland et al., 2012; Zhang et al., 2018; Weckesser et al., 2019) and low level to no correlation between biological measures and questionnaires (Vanaelst et al., 2012; Stalder et al., 2017; Prado-Gascó et al., 2019). Some scholars suggest log-transformation of cortisol values to better approximate a Gaussian distribution of cortisol values across subjects (Lenow et al., 2017; Lempert et al., 2018a). Indeed, we decided it is reasonable to modify all of our cortisol values based on the distributions before and after log-transformation (Figure S5, SI).

From the correlation analysis where we combined all stress measures, cortisol levels were not associated with questionnaire scores. Overall, we found more significant correlations within the stress measure groups (which may validate stress measures per se), compared to across stress measure groups (Figure 3B). Still, as expected aggregated salivary cortisol (ls123) was positively correlated with two hair cortisol samples (ls123 vs. lh2: Pearson *r* = 0.38, *p* = 0.024; ls123 vs. lh3: Pearson *r* = 0.35, *p* = 0.039; as in Weckesser et al. (Weckesser et al., 2019)). With the multiple comparison correction (8 comparisons per stress variable: *α*-level 0.05/8) only positive correlations between lh2 and lh3 (Pearson *r* = 0.46, *p* = 0.005) and PSS and BEPSI questionnaires (Pearson *r* = 0.78, *p* < 0.001) remained significant. Stronger relations within the stress measure groups were further confirmed with a principle component analysis (PCA: Figure S6 & Table S4, SI): the second principle component along the nine stress-vectors (all stress variables) distinguished between human bio samples (hair and saliva) and questionnaires, whereas the third principle component distinguished between hair and saliva stress measures. By and large, cortisol values for saliva (ng/mL) and hair (pg/mg) samples stayed within the same ranges, respectively, across experimental sessions (Figure 3C). According to permutation tests hair samples were not significantly different, while log(cortisol) for saliva session 1 sample was significantly higher than that for the other sessions (e.g., log(cortisol) from saliva sample: *M*_*s*1_ = − 0.047 & *M*_*s*3_ = − 0.330, *p* = 0.011).

Literature suggests that because gender is associated with neuroendocrinological substrates (e.g., cortisol, Takahashi, 2004; Barel et al., 2017), the relationships between stress-related hormones and decision making should control for gender. Moreover, there are studies that only include males due to reported menstrual and contraceptive effects on cortisol levels (Lempert et al., 2012; Stalder et al., 2017). We did not find effects of gender for all stress variables using permutation tests (*p* > 0.09). Also, by visually assessing the differences through the first three principle components and respective 2D planes we found that the gender ellipses as well as the data were highly overlapping (Figure S6, SI).

### 2.3 Stress variables were selectively predictive of seconds discount factor

It is important to distinguish between the effects of endogenous variation in stress on within-subject and on between-subject variability in choice preferences. Both can be explored using the stress measures and the economic tasks in this study (Figure 1).

First, given repeated economic tasks and biological measures corresponding to the experimental sessions/months we could check whether within-subject changes in behavior were related to changes in stress levels. In particular, we wanted to explore subtle deviations from the base level (given that discount factors and cortisol measures were stable on average in Figures 2D & 3C). Second, considering that the ranges of stress measures in our study reflected that of healthy population (Table S3, SI) we explored whether stress measures per session/month shared some intersubject (between-subject) variance with the economic behavior. In other words, using a combination of stress measures over time can we predict subjects’ behavior in delay-discounting tasks?

Correlational analysis revealed a significant positive correlation between the change in the seconds delay discount factor (log(*k*_*SDD*2_) − log(*k*_*SDD*1_)) and hair cortisol change (Δlh3 = lh3 - lh1, Pearson *r* = 0.36, *p* = 0.033, Figure 4A, no longer significant if we used the Bonferroni multiple comparisons correction; other correlations in Table S5, SI). Then we ran a conventional set of regression models (including Lasso, Ridge, and Elastic Net regressions) with 5-fold cross validation and standardized data (Materials and Methods). Ridge regression outperformed other models and the only model that had the coefficient of determination (*R*^2^) bigger than 0.2 was predicting the change in the seconds delay discount factor (Table 1 & Figure 4B-D). We used the SHapley Additive exPlanations (SHAP) to interpret the model predictions (Materials and Methods).

**Table 1.** Ridge Regression Comparison.

| **within-subject**<br>N=36 | $\log(k_{SDD2})$<br>$-\log(k_{SDD1})$ | $\log(k_{DDD2})$<br>$-\log(k_{DDD1})$ | $\log(k_{DDD3})$<br>$-\log(k_{DDD1})$ | $\log(k_{DDD3})$<br>$-\log(k_{DDD2})$ | |
| --- | --- | --- | --- | --- | --- |
| $R^2$ | 0.25 | 0.13 | 0.06 | 0.10 | |
| **between-subject**<br>N=34 | $\log(k_{SDD1})$ | $\log(k_{SDD2})$ | $\log(k_{DDD1})$ | $\log(k_{DDD2})$ | $\log(k_{DDD3})$ |
| $R^2$ | 0.50 | 0.24 | 0.18 | 0.06 | 0.07 |

*Note.* Each column identified the dependent variable (the target). The within-subject models included six independent variables (features): $\Delta ls2$ , $\Delta ls3$ , $\Delta ls32$ , $\Delta lh2$ , $\Delta lh3$ , $\Delta lh32$ . The between-subject models included nine independent variables (features): pss, bepsi, lcu, ls1, ls2, ls3, lh1, lh2, lh3. All variables were standardized before their use in the models. Ridge regressions were set up with the use of 5-fold cross validation and a search through parameter space for the best $\alpha$ from (0.001, 0.01, 0.1, 1, the penalty term in the Ridge regression). In Figure 5 the plots showed the results of predicting the seconds discount factor averaged across sessions (i.e., the dependent variable $\log(k_{SDD}) = (\log(k_{SDD1}) + \log(k_{SDD2}))/2$ ). This model had $R^2 = 0.38$ .

**Figure 4.**
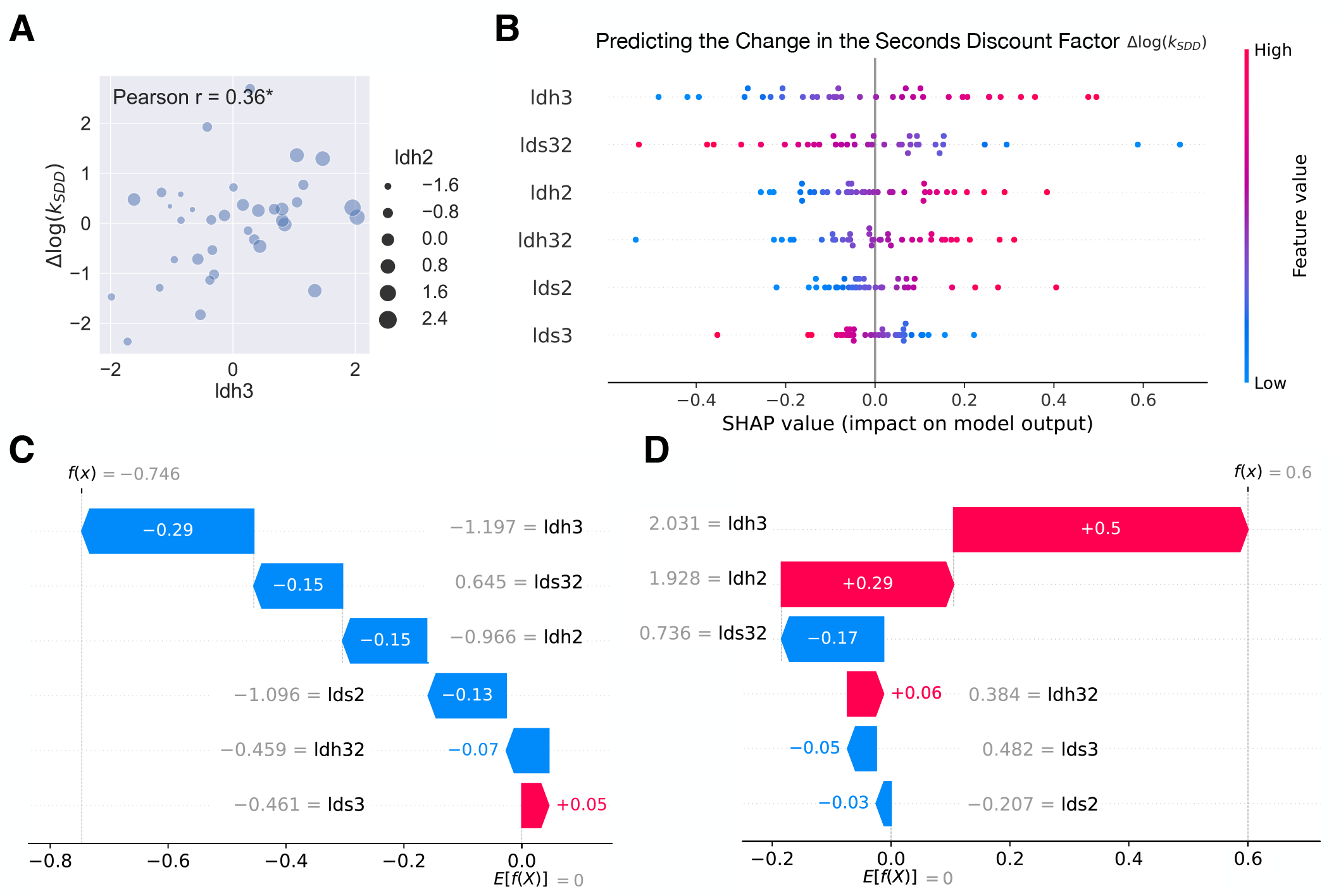
Within-subject analysis. (A) Scatterplot of change in hair cortisol (Δlh3, x-axis) and change in seconds discount factor (Δ log(*k*_*SDD*_), y-axis). The size of the dots was aligned with Δlh2. Pearson *r* was reported on the plot (*r* = 0.36, *p* = 0.033; * for *p* < 0.05). (B) SHAP summary plot (beeswarm) that explained the within-subject Ridge regression predictions of the change in the seconds delay discount factor (log(*k*_*SDD*2_) − log(*k*_*SDD*1_)). The independent variables (features) were displayed in descending order by importance. Each subject was represented by a single dot on each variable row. The x position of the dot was determined by the SHAP value of that feature, i.e. showing whether the effect of that value was associated with a higher or lower prediction. Color was used to display the original value of a feature. (C) & (D) Examples of two extreme subjects’ predictions. The waterfall plot explained predictions for individual subjects, i.e. it showed how the positive (red) or negative (blue) contribution of each feature moved the value from the expected value of the change in the seconds delay discounting across sessions to the value of this subject’s prediction. Numbers on the arrows or in color signified individual SHAP values, numbers in gray were the standardized values of the stress variables of the subject under consideration.

According to the SHAP summary plot (Figure 4B) first three features in the order of importance were Δlh3, Δls32 = ls3 - ls2, Δlh2 = lh2 - lh1. Having seen the correlation before, not surprisingly the hair cortisol change had the biggest positive impact on the prediction of the change in seconds discount factor (consistent with our hypothesis). We decided to unpack the SHAP summary further by focusing on the two example subjects located at the extremes: same three stress variables appear as top predictors. For one of them the 2 standard deviations increase in stress level across months (Δlh3 = 2.031) was the single main contributor to the prediction of increased impulsivity (by 0.5 standard deviations, Figure 4D) in the seconds delay task, for the second one the 1 standard deviation decrease in stress level across months (Δlh3 = -1.197) was the main contributor to the prediction of increased patience (by 0.29 standard deviations, Figure 4C) in the seconds delay task.

In the between-subject analysis, average seconds delay discount factor correlated with the life change units (log(*k*_*SDD*_) vs lcu: Pearson *r* = 0.38, *p* = 0.029), and the seconds delay discount factor in the session 1 correlated with the second principle component (Pearson *r* = 0.52, *p* = 0.002, remaining significant after the Bonferroni multiple comparisons correction: *α*-level 0.05*/*13; PC2 along the nine stress-vectors: three saliva, three hair, and three questionnaires represented a direction that separated human bio samples and self-reported measures in Figure S6, SI; further PCA details and other correlations in SI).

As in the within-subject analysis, Ridge regression outperformed other regressions (Lasso and Elastic Net, all fit with 5-fold cross validation and standardized data, Materials and Methods) with the coefficient of determination (*R*^2^) bigger than 0.2 for models predicting the seconds delay discount factor (Table 1 & Figure 5C). SHAP for the between-subject analysis again highlighted the importance of hair stress measure with lh3, representing the stress level during the month of the sample collection, being the key negative predictor of the discount factor. Also, the stress level the month before the collection (lh2) provided one of the biggest positive contributions to the prediction of the seconds discount factor. Only the latter finding was expected (given our hypothesis derived from the literature) and intuitive: the higher the stress level the higher the seconds discount factor relative to the average (i.e., more impulsive, Figure 5C). Another important feature in the between-subject regression (and in the two separate regressions with the seconds discount factor in sessions 1 & 2 as dependent variable) is the life change units (lcu) that gave the biggest positive contribution to the seconds discount factor: the more traumatic experiences a subject experienced (higher lcu) the more they discounted in seconds (higher discount factor was associated with increased impulsivity, again as per our hypothesis, Figure 5A).

**Figure 5.**
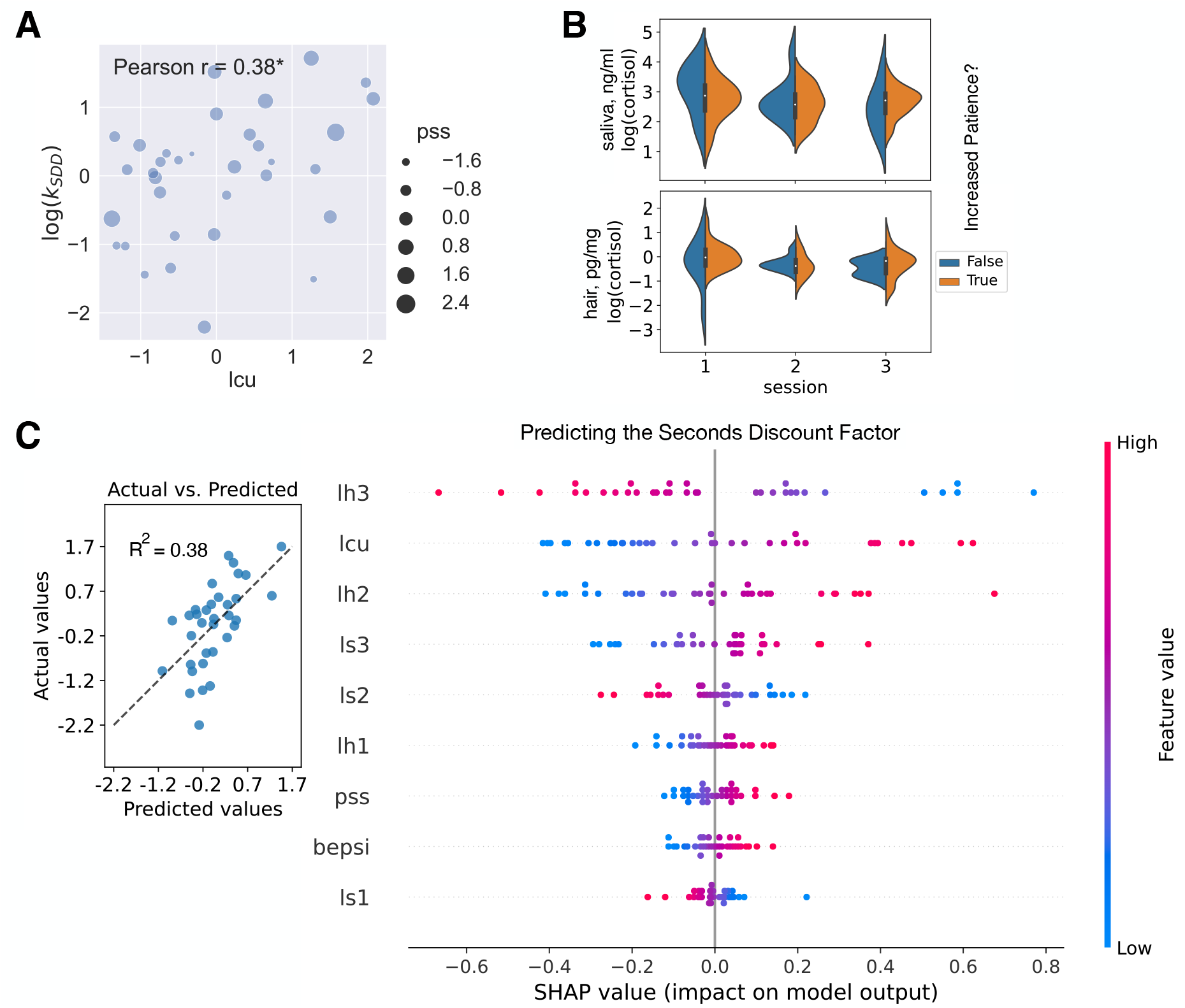
Between-subject analysis. (A) Scatterplot of life change units (lcu, x-axis) and seconds discount factor averaged across sessions (log(*k*_*SDD*_), y-axis). The size of the dots reflected another self-reported stress measure, perceived stress scale (pss). Pearson *r* was reported on the plot (*r* = 0.38, *p* = 0.029; * for *p* < 0.05). (B) Violin plot visualizing no significant differences between log(cortisol) in saliva and hair samples between subjects who increased their patience in the day delay task in session 3 and those who did not. (C) Cross-validated predictions from the between-subject Ridge regression with the following dependent variable: log(*k*_*SDD*_) = (log(*k*_*SDD*1_) + log(*k*_*SDD*2_))*/*2. In the left subplot the predicted values (x-axis) were plotted against the actual values (y-axis). The coefficient of determination was shown on the plot. The dashed line was the best fit line. In the right subplot, the SHAP summary plot (beeswarm) explained the between-subject Ridge regression predictions. The independent variables (features) were displayed in descending order by importance. Each subject was represented by a single dot on each variable row. The x position of the dot was determined by the SHAP value of that feature, i.e. showing whether the effect of that value was associated with a higher or lower prediction. Color was used to display the original value of a feature.

Finally, we checked whether we can classify subjects who increased their patience over the gained experience with the day delay task given their stress profile. There were 27 such subjects, as opposed to 7 subjects whose discount coefficient stayed the same level or increased (decrease in patience). According to three basic classifiers (logistic regression, decision tree, and random forest, with the use of 5-fold stratified cross validation; Materials and Methods) the stress variables were not helpful to dissociate between increased and decreased patience (with overlapping distributions of stress supporting this result in Figure 5B). For example, the logistic regression was able to predict the majority class (increase in patience) well with 79.5% accuracy, but did not predict minority class at all, leaving the balanced accuracy at the chance level (50%) and area under curve even below 50%. Therefore, temporal context, or the fact that the day delay task was paired with the week delay task in session 3, could be the answer.

## 3 Discussion

In our study, biological stress measures were moderately correlated between each other as seen in the literature before, further validating the stress measurement system (Vanaelst et al., 2012; van Holland et al., 2012; Zhang et al., 2018; Weckesser et al., 2019). However, overall we found more significant relationships within stress modalities, rather than across them (also in PCA analysis, SI). This gives more evidence that self-reported stress, salivary cortisol, and hair cortisol might affect decision making by operating via distinct channels.

The longitudinal within-subject design allowed us to find effects of biological stress markers and their changes over time both on within-subject and between-subject behavioral variability. The strongest result we found for the within-subject variability was that the increases in the levels of cortisol were associated with increased impulsivity in the seconds delay task across experimental sessions, which was consistent with our hypothesis. Still, the intuition behind the positive effect of change in hair cortisol (Δlh3) but negative effect of change in saliva cortisol (Δls32) could not be grasped. The timing of comparison was slightly different: a month delta for the hair and two weeks for the saliva, but we would not speculate about this further. In the between-subject analysis, although hair stress level appeared among important regressors to predict subjects’ time preference over seconds, the direction of influence for stress in the month of sample collection (lh3) compared to stress in the month prior to the collection (lh2) was directly opposite. Only life change units contributed to the prediction in the expected manner: more stress linked to higher impulsivity in seconds as per our hypothesis derived from previous studies. Additionally, models that captured behavior from the seconds delay discounting task did relatively better than those using a traditional delay discounting task. The seconds delay discounting task was the only task where all trials contributed to the total profit and the total waiting time. In other words, endogenous stress was more prone to influence behavior in a ‘real’ experiential task rather than in a task with potential outcomes determined by a random draw in the end of the experimental session.

Previous literature accepts both linear and curvilinear relationships between changes in cortisol as a physiological response to stress and behavior (Goldfarb et al., 2017; Maier et al., 2015; Luksys and Sandi, 2011; Lempert et al., 2018a). We did not find any curvilinear relationships between stress variables and model-free behavioral measures in a separate analysis (SI). This might be due to the fact that tests for curvilinear relationships included only a small portion of participants, those for whom there was an increase in biological stress level compared to the baseline. In fact, SHAP values would not capture nonlinear relationships in our regression analysis and would assume that the model is additive, i.e. that the contribution of each feature is independent of the other features, an assumption that might not hold. Although having nice properties, including visualizations, the explanation we provided for the Ridge regression model results using SHAP values is by no means causal: SHAP values show how each independent variable contributed to a prediction, they do not tell how the independent variables contributed to the dependent variable.

Among self-reported stress proxies the life change units were positively correlated to the seconds discount factor and negatively correlated to the proportion of later choices in the seconds delay task. It is worth noting that life change unit is a measure corresponding to real traumatic experience of subjects in a particular event, compared to two other questionnaires that asked for feelings about (The Brief Encounter Psychosocial Instrument) or subjective appraisal of (Perceived Stress Scale) particular life events or situations. Previous scholarship did find a robust relationship between stressful life events and increased negative health outcomes, however, just a few studies investigate life change units’ relationship with decision making (Wang et al., 2020). Although we found that those with more traumatic experiences discount more, as expected, it might be only true for those with lower cognitive speed (Friedel, 2017). The lower levels of cognitive speed could potentially worsen conversion from seconds to days when subjects are presented with choices in the seconds delay task or provide less capacity for waiting in real time. We did not use any measures of cognitive speed in this study to test this explanation.

In our study design, there were several limitations that need to be acknowledged. First, delay-discounting tasks were not measured an equal number of times, which may have affected our results. This was due to both the desire to replicate previous behavioral results (Lukinova et al., 2019) in a general population group and to test more economic tasks within the same subjects. Second, questionnaires were not carried out repeatedly. Having questionnaires that reflect the self-reported stress in the within-subject regression might indicate new relationships between subtle changes in stress and behavior. Finally, our study suffered from a small sample size and high attrition rate (17%) making our sample size too low for a predominantly correlational study, therefore resulting in insufficient statistical power to detect significant effects. The limitations listed should be taken into consideration when interpreting the results of this study.

Our study did not induce stress in participants but rather examined cortisol levels, their changes, and fluctuations from the baseline, which may be due to individual responses to the environment. This is rather new. Previously in control groups random fluctuations of cortisol did not yield the same behavioral patterns (Lempert et al., 2018a). However, the majority of previous studies were using saliva measure (Takahashi, 2004; Linz et al., 2018). Most of our significant relationships between stress and behavior included the hair cortisol measure. We conclude that for subtle changes in behavior due to random cortisol fluctuations hair stress measure might be more reliable.

Measuring and administering hormones is a complex task, as hormone levels vary throughout the day and month. For example, stressors that are uncontrollable tend to produce a high overall level of daily cortisol release, compared to controllable stressors - higher-than-normal morning cortisol (Miller et al., 2007). As a result, many studies involving hormones and decision-making produce conflicting results or fail to be replicated and, therefore, are not able to advance the current state of the discipline. We should reiterate that we are pioneers to combine self-reported stress, biological stress, and delay-discounting tasks across time horizons. Most of our significant relationships were between endogenous variation in hair cortisol and the seconds delay task that were never studied together. Some of our main results support the hypothesis in the literature that increased stress leads to steeper discounting. Still, we found that endogenous subclinical variation in cortisol only weakly correlated with economic preferences. The strongest links we found were that within-subject increases in stress level led to increases in discounting and higher levels of stress between-subject were related to higher discounting, both in the seconds delay task.

Additional multi-task and multi-stress studies with higher *N* are required to establish effects discovered in this paper. An accurate understanding of the biological processes that lead people to discount future payoffs is still tenuous at best. Unraveling mechanisms of how chronic stress spreads unfavorable effects to behavior is of critical importance to policymakers and of strong interest to the general public.

## 4 Materials and Methods

### 4.1 Participants

In this study, we recruited 44 participants total (26 female, 17 male, 1 other) including 15 participants from New York University (NYU) Shanghai staff and 29 participants from Chinese working population around NYU Shanghai and East China Normal University (ECNU) campuses. We pre-screened subjects before recruitment and excluded smokers, drinkers, and subjects taking medicine, or suffering from acute or chronic hormonal dysregulation. Three participants withdrew after the first session and the rest (41: 23 female, 17 male, 1 other, *M*_*age*_ = 30) completed the whole experiment. All of our participants shared a similar socioeconomic status (Figure S1A, SI). 31 participants (out of 41) had no siblings, 22 (out of 41) were not married and 32 (out of 41) did not have any chronic illnesses.

The recruitment and experimental sessions for the study were completed between 2nd April and 20th October, 2018. The study was approved by the Institutional Review Board (IRB) of NYU Shanghai. All subjects gave informed consent (written) before participation in the study. All research was performed in accordance with relevant guidelines and regulations.

### 4.2 Experimental Tasks

There were three behavioral task sessions in total. They were scheduled approximately every two weeks and took place in the NYU Shanghai Behavioral and Experimental Economics Laboratory, or the experimental room in the Geo Building at ECNU. Each session consisted of a two-alternative choice task. In the first two sessions, participants made their choices between a later option (delay in seconds and days) and an immediate option (now and today). The last session involved a delay-discounting task comprising choices between a later option (delay in days and weeks) and an immediate option (today). Participants received a 30 CNY ($4.3 USD) per session participation fee as well as up to an additional 70 CNY ($9.9 USD) per session based on their individual performance in the tasks.

We followed the same procedure for each delay-discounting task that was established in our previous works (Lukinova et al., 2019; Lukinova and Erlich, 2021). The main difference was that both instruction and the screen stimuli were presented in Chinese. During the first two sessions, subjects experienced an alternating set of four tasks with the order counter-balanced. Therefore, subjects could experience one of the following sequences: SDD-DDD-SDD-DDD or DDD-SDD-DDD-SDD. During the third session, subjects experienced two tasks with the order counter-balanced, i.e., DDD-WDD or WDD-DDD.

In each trial of SDD, subjects made a decision between the ‘now’ and the ‘in *n* seconds’ options (Figure 2A). If subjects chose a delayed option, then they had to experience the delay right after choice, e.g., they had to watch the clock for 30 seconds until that screen disappeared and they could continue to the next trial. The ‘now’ reward was fixed at 4 coins, the ‘later’ rewards used in the trials were 1, 2, 5, 8, 10 coins (5 in total) combined with delay to rewards: 3, 6.5, 14, 30, and 64 seconds (5 in total). The two ‘smaller-later’ options (1, 2) made up 10% of later options. Subjects accumulated coins which were transferred in the end of the task to CNY following the exchange rate in the instruction.

In contrast, for other delay-discounting tasks (with delays in days or in weeks) subjects made a decision between the ‘today’ and the ‘in *n* days/weeks’ options (Figure 2A), saw their choice confirmation and then proceeded to the next trial. The coins were not accumulated. The ‘today’ reward was fixed at 4 coins, the ‘later’ rewards used in the trials were the same as in SDD with the same values of delays to rewards in days for DDD and in weeks for WDD. At the end of the session, a single trial was selected randomly, and the coins earned in that trial were transferred to CNY following the exchange rate (much bigger than the one used in SDD) to determine the payment. If in the randomly selected trial subject chose the later option, the payment was transferred only after several days or weeks delay.

### 4.3 Stress Measures

We used three questionnaires to measure perceived stress. The 10-item version of the Perceived Stress Scale (PSS, Cohen et al., 1983) was used to assess an individual’s subjective appraisal of particular life events/situations as being unpredictable, uncontrollable, and/or overloaded. On a 5-point Likert scale ranging from 0 ‘never’ to 4 ‘very often’, participants rated how often in the previous month they felt or thought as described in 10 examples. The Brief Encounter Psychosocial Instrument (BEPSI, Frank and Zyzanski, 1988) usually consists of 6 questions. We omitted the first open-ended question and used the rest: 5 closed-ended questions. Subjects were asked to respond yes or no to each item and if yes, to rate the impact of these stressors on a scale of 1 to 10. The Social Readjustment Rating Scale (SRRS, Holmes and Rahe, 1967) was used for identifying major stressful life events. Each one of the 43 stressful life events was awarded a Life Change Unit (LCU) depending on how traumatic it was felt to be by a large sample of participants. The total then was calculated by adding all LCUs for the events a subject experienced (‘lcu’ variable). Thus, in all questionnaires higher scores were associated with higher levels of perceived stress. All questionnaires were administered in Chinese.

In order to obtain another measure of participant’s stress level, in the beginning of each session, we collected a saliva sample from each participant. In healthy individuals, cortisol levels peak in the early morning, and gradually decrease thereafter. To help overcome this challenge, most studies obtain multiple human specimen samples from the time of waking up until sleep or collect samples at the same time for several days. For each participant, the collection time remained the same during three sessions (as in Pruessner et al., 2003; González-Cabrera et al., 2014). Therefore, we could compare salivary cortisol levels to the baseline (session 1) and attribute these deltas to the changes from the base cortisol level. Saliva samples were collected using the Salivette systems. The Salivette systems (Sarstedt, Shanghai) came with synthetic swabs. Subjects were asked to refrain from eating, drinking and/or brushing teeth for at least 1 hour before sampling and to rinse their mouth 10 minutes before arriving to the experimental facility. Before each experimental session, experimenter asked a subject to place the swab in the mouth, chew it for 1 min and then transfer it into the tube. All subjects had three saliva samples of necessary quality. The samples were stored on ice till the end of the experimental day, then, were centrifuged and stored at -20 °C until analysis at NYU Shanghai Molecular Biology Lab. On the day of analysis, we measured salivary cortisol by enzyme-linked immunosorbent assay (ELISA, IBL International, Germany). Saliva samples were assayed in duplicate (40 * 2 samples per ELISA kit) and measured according to the instructions of the kits. The controls were within expected bounds.

Two weeks after all three experimental sessions were completed by subjects, we also collected their hair samples to measure the levels of chronic stress. 36 subjects returned to provide hair samples. Hair samples were collected by cutting and merging at the cut two strands of hair (with the total thickness of a toothpick each) from posterior vertex area. All samples were stored in aluminum foil at room temperature. LC-MS/MS (liquid chromatography with tandem mass spectrometry) atmospheric pressure chemical ionization (APCI, MRM mode) was used to analyze cortisol in hair samples. The analysis was performed by Dr. Huihua Deng’s lab at Southeast University in China using their standard approach (Chen et al., 2013). Hair strands longer than 3 cm were cut as close to the scalp as possible and segmented into three 1cm-long parts. All segmented samples were labelled as SXxx-01-SXxx-03 from bottom (closest to the scalp) to top that later became lh3 - lh1 variables, respectively. In total 36*3 = 108 samples were analyzed. The recovery rate was within 90-105%, and intra-day and inter-day coefficients of variation were below 10%, which satisfied measurement requirement.

### 4.4 Analysis

#### 4.4.1 Fitting Economic Choices

Subjects’ discount factors were estimated by fitting their choices with a Bayesian hierarchical model (BHM) of hyperbolic discounting with decision noise and a reward scaling parameter, ‘rews’ that scaled the delayed reward per unit. Bayesian hierarchical modeling allows for pooling data across subjects, while differentiating between two delay-discounting tasks per session and recognizing subject differences. Among its advantages is also the ability to estimate posterior distributions, rather than getting only the point estimates of the parameters. We did not compare different functional forms and followed our previous work by using the softmax-hyperbolic model (Lukinova et al., 2019). Compared to the exponential model (Samuelson, 1937) of time discounting that has a straightforward economic meaning - a constant probability of loss of reward per waiting time, the hyperbolic model (Mazur, 1987) according to the scholarship more accurately describes how individuals discount future rewards. At the same time a shift-invariant softmax rule is commonly used in the field to transform the subjective utilities of the sooner and later offers into a probability of choosing the later offer.

The main ‘reward scale’ model (*M*_6*p*,3*s*_) had 6 population level parameters (log(*k*), decision-noise, log(*τ*), and reward scale parameter, rews, for each of the two delay-discounting tasks per session) and 3 parameters per subject: e.g., for session 1 log(*k*_*SDD*1_), log(*k*_*DDD*1_) and log(*τ*).

~~~
n_ chose _ later | trials(n_ trials) ∼ (exp (rews)* later_ reward / (1 + exp (logk) * delay) - sooner
      _ reward) / exp (logtau),
logk ∼ unit + (unit|subjid),
logtau ∼ unit + (1| subjid),
rews ∼ unit
~~~

where `later_reward` was the later reward, `sooner_reward` was the sooner reward; `logk` was the natural logarithm of the discounting parameter *k* and `logtau` (log(*τ*)) was the natural logarithm of the decision noise; in the binomial specification the data was grouped and summarized by distinct trial types, where `n_chose_later` was the count of choices when the later reward was selected and `n_trials` was the number of trials for a particular trial type.

Based on the 10-fold cross-validation criteria the main model was preferred to the models with reduced numbers of parameters (Table S1, SI). The main model was estimated after conversion of all delays to days (resulted in the same fits as the fits in the units of the task) with the ‘brms’ package in R (Bürkner, 2017) that allows to do BHM of nonlinear multilevel models in Stan (Carpenter et al., 2016) with the standard R formula syntax.

#### 4.4.2 Making Comparisons

The permutation tests of differences between the means of two groups were done by shuffling the group label and computing the mean between the shuffled groups 10000 times. This generated a null distribution which was used to estimate the probability of observing the true difference between groups (bootmean in https://github.com/erlichlab/elutils). The visualizations of the correlation matrices were done via R package ‘corrplot’ (Friendly, 2002). Whenever multiple hypotheses were tested at the same time (e.g., multiple correlations were run between discount factor and the stress variables) we used the Bonferroni multiple comparison correction with *α*-level being *α/n*, where *n* was the number of hypotheses.

#### 4.4.3 Predictive Modeling

We used Lasso, Ridge, and Elastic Net cross-validated (5-fold) regressions with standardized data to predict discount factors and their change using stress variables (within-subject and between-subject analyses). In particular, ‘sklearn’ Python library classes ‘sklearn.linear model.LassoCV’, ‘RidgeCV’, ‘ElasticNetCV’ were used, respectively. We tried an array of *α* values (penalties: 0.001, 0.01, 0.1, 1). Once the best penalty parameter was found through cross validation, the model was fit again using the entire training set. We chose these models due to their ability to address issues of overfitting and multicollinearity. They do so by adding a penalty term to the cost function that shrinks the magnitude of the estimated coefficients, leading to better generalization performance and more robust predictions. Ridge regression (where regularization is given by the l2-norm) outperformed other models by the coefficient of determination and can be especially relevant for our data given correlations among input features and a small dataset.

The SHapley Additive exPlanations (SHAP, using Python library ‘shap’) were imperative to explain predictions of our best Ridge regressions. Following Lundberg and Lee (2017) we imagined that regression model inputs are players and model predictions are players’ payouts, then SHAP is able to provide the contribution of each player to the game. Not only SHAP values combined can show which features are important and the direction of correlation with the prediction (Figure 4B), but also for each individual prediction (e.g., prediction of the change in subject’s seconds discount factor) individual SHAP values detail the contribution of each feature and show how each feature moves the value from the expected value of output feature (dependent variable) to this particular subject’s prediction (e.g., for two extreme subjects Figure 4C & D).

Finally, in order to predict the classes of subjects who increased/decreased their patience in the delay task over the sessions we used three commonly used classifiers, such as logistic regression, decision tree, and random forest (all through Python library ‘sklearn’). The ‘tree’ non-parametric models were considered to allow for complex nonlinear relationships between input features and the output. We implemented the 5-fold stratified cross validation and compared the models using balanced accuracy and the area under the receiver operating characteristic curve (AUC ROC) due to the class imbalance (out of 34 subjects 27 increased in patience and only 7 subjects stayed at the same level or decreased in patience). Stratified cross validation helped us to ensure that the model was trained and tested on representative subsets of the data, reducing the risk of overfitting and improving the generalization performance of the model. The logistic regression outperformed other models in the ability to predict the majority class (by accuracy), but was not able to predict the minority class (checked via confusion matrices), therefore, resulting in predictions at or below the chance level (50%).

### 4.5 Software

The code for the delay-discounting task was written in Python using the ‘PsychoPy’ toolbox (version 1.83.04; Peirce, 2007) available at https://www.github.com/erlichlab/gpstress/src/task. All analyses and statistics were performed either in Matlab (version 9.3, or higher, The Mathworks, MA), in R (version 3.6.1, or higher, R Foundation for Statistical Computing, Vienna, Austria), or in Python (version 3.9.13).

### 4.6 Data Availability

Software for running the task, as well as the data and analysis code for regenerating our main results and figures are available as a Zenodo release (https://doi.org/10.5281/zenodo.5513429) of a GitHub repository (https://github.com/erlichlab/gpstress).

## Author Contributions

All authors contributed to the study design, interpretation of the results and writing the manuscript. Data collection and analyses were performed by E. Lukinova under the supervision of J. Erlich.

## Acknowledgments

We acknowledge an outstanding NYU Shanghai undergraduate who helped us to start this project: Yuyue Wang. We acknowledge Brianna Fu from NYU, Danielle John from CUNY Hunter College and Shengjie Xu from NYU Shanghai who helped us to recruit Chinese general population participants and run sessions. We thank NYU Shanghai undergraduate students Xirui Zhao and Muzi Du who helped us to run ELISA on saliva samples. We acknowledge Prof. Wenshu Li who allowed and supported our use of NYU Shanghai Molecular Biology Lab for running ELISA. Funding was provided by National Science Foundation of China (NSFC-31750110461) International Young Scholar Grant and Shanghai Eastern Scholar Program (Shanghai Municipal Education Commission) Talents Young Scholar Grant to E.L.

## References

David Åkerlund, Bart HH Golsteyn, Hans Grönqvist, and Lena Lindahl. Time discounting and criminal behavior. Proceedings of the National Academy of Sciences, page 201522445, 2016.

American Psychological Association et al. Stress in america 2020: A national mental health crisis. 2020, 2021.

Efrat Barel, Shosh Shahrabani, and Orna Tzischinsky. Sex Hormone/Cortisol Ratios Differentially Modulate Risk-Taking in Men and Women. Evolutionary Psychology, 15(1):147470491769733, January 2017. ISSN 1474-7049, 1474-7049. doi: 10.1177/1474704917697333. URL http://journals.sagepub.com/doi/10.1177/1474704917697333.

Paul-Christian Bürkner. brms: An R Package for Bayesian Multilevel Models Using Stan. Journal of Statistical Software, 80(1):1–28, 2017. doi: 10.18637/jss.v080.i01.

Bob Carpenter, Andrew Gelman, Matt Hoffman, Daniel Lee, Ben Goodrich, Michael Betancourt, Michael A Brubaker, Jiqiang Guo, Peter Li, and Allen Riddell. Stan: A probabilistic programming language. Journal of Statistical Software, 20:1–37, 2016.

Smarandita Ceccato, Brigitte M Kudielka, and Christiane Schwieren. Increased risk taking in relation to chronic stress in adults. Frontiers in psychology, 6, 2015.

Zheng Chen, Jifeng Li, Jing Zhang, Xue Xing, Wei Gao, Zuhong Lu, and Huihua Deng. Simultaneous determination of hair cortisol, cortisone and DHEAS with liquid chromatography–electrospray ionization-tandem mass spectrometry in negative mode. Journal of Chromatography B, 929:187–194, June 2013. ISSN 15700232. doi: 10.1016/j.jchromb.2013.04.026. URL https://linkinghub.elsevier.com/retrieve/pii/S1570023213002365.

Sheldon Cohen, Tom Kamarck, and Robin Mermelstein. A Global Measure of Perceived Stress. Journal of Health and Social Behavior, 24(4):385, December 1983. ISSN 00221465. doi: 10.2307/2136404. URL http://www.jstor.org/stable/2136404?origin=crossref.

Alain Cohn, Jan Engelmann, Ernst Fehr, and Michel André Maréchal. Evidence for countercyclical risk aversion: an experiment with financial professionals. The American Economic Review, 105(2):860–885, 2015.

Andrea Dietrich, Johan Ormel, Jan K Buitelaar, Frank C Verhulst, Pieter J Hoekstra, and Catharina A Hartman. Cortisol in the morning and dimensions of anxiety, depression, and aggression in children from a general population and clinic-referred cohort: an integrated analysis. The TRAILS study. Psychoneuroendocrinology, 38(8):1281–1298, 2013.

Carlo Faravelli and Stefano Pallanti. Recent life events and panic disorder. Am J Psychiatry, 146(5):622–6, 1989.

Scott H Frank and Stephen J Zyzanski. Stress in the clinical setting: the Brief Encounter Psychosocial Instrument. The Journal of family practice, 26(5):533–9, June 1988.

Eva Friedel. How Accumulated Real Life Stress Experience and Cognitive Speed Interact on Decision-Making Processes. Frontiers in Human Neuroscience, 11, 2017. ISSN 16625161. doi: 10.3389/fnhum.2017.00302. URL http://journal.frontiersin.org/article/10.3389/fnhum.2017.00302/full.

Michael Friendly. Corrgrams: Exploratory Displays for Correlation Matrices. The American Statistician, 56(4):316–324, November 2002. ISSN 0003-1305, 1537-2731. doi: 10.1198/000313002533. URL http://www.tandfonline.com/doi/abs/10.1198/000313002533.

Arline T. Geronimus, Margaret T. Hicken, Jay A. Pearson, Sarah J. Seashols, Kelly L. Brown, and Tracey Dawson Cruz. Do US Black Women Experience Stress-Related Accelerated Biological Aging?: A Novel Theory and First Population-Based Test of Black-White Differences in Telomere Length. Human Nature, 21(1):19–38, March 2010. ISSN 1045-6767, 1936-4776. doi: 10.1007/s12110-010-9078-0. URL http://link.springer.com/10.1007/s12110-010-9078-0.

Elizabeth V. Goldfarb, Monja I. Froböse, Roshan Cools, and Elizabeth A. Phelps. Stress and cognitive flexibility: Cortisol increases are associated with enhanced updating but impaired switching. Journal of Cognitive Neuroscience, 29(1):14–24, January 2017. ISSN 0898-929X, 1530-8898. doi: 10.1162/jocn_a_01029. URL https://www.mitpressjournals.org/doi/abs/10.1162/jocn_a_01029.

Bart HH Golsteyn, Hans Grönqvist, and Lena Lindahl. Adolescent time preferences predict lifetime outcomes. The Economic Journal, 124(580):F739–F761, 2014.

J González-Cabrera, M Fernández-Prada, C Iribar-Ibabe, and JM Peinado. Acute and chronic stress increase salivary cortisol: a study in the real-life setting of a national examination undertaken by medical graduates. Stress, 17(2):149–156, 2014.

Holly C Gooding, Carly E Milliren, S Bryn Austin, Margaret A Sheridan, and Katie A McLaughlin. Child abuse, resting blood pressure, and blood pressure reactivity to psychosocial stress. Journal of pediatric psychology, page jsv040, 2015.

Lisa A Goodman, Carole Corcoran, Kiban Turner, Nicole Yuan, and Bonnie L Green. Assessing traumatic event exposure: General issues and preliminary findings for the Stressful Life Events Screening Questionnaire. Journal of traumatic stress, 11(3):521–542, 1998.

R. Gow, S. Thomson, M. Rieder, S. Van Uum, and G. Koren. An assessment of cortisol analysis in hair and its clinical applications. Forensic Science International, 196(1-3):32–37, March 2010. ISSN 03790738. doi: 10.1016/j.forsciint.2009.12.040. URL https://linkinghub.elsevier.com/retrieve/pii/S0379073809005416.

Constance Hammen. Stress and depression. Annu. Rev. Clin. Psychol., 1:293–319, 2005.

Allison G Harvey, Charlie Jones, and D Anne Schmidt. Sleep and posttraumatic stress disorder: a review. Clinical Psychology Review, 23(3):377–407, 2003.

Johannes Haushofer and Ernst Fehr. On the psychology of poverty. Science, 344(6186):862–867, May 2014. doi: 10.1126/science.1232491. URL http://www.sciencemag.org/cgi/content/abstract/344/6186/862.00000.

Dirk H Hellhammer, Stefan Wüst, and Brigitte M Kudielka. Salivary cortisol as a biomarker in stress research. Psychoneuroendocrinology, 34(2):163–171, 2009.

Thomas H Holmes and Richard H Rahe. The social readjustment rating scale. Journal of psychosomatic research, 11(2):213–218, 1967.

Lisa Huang, Amir Rattner, Han Liu, and Jeremy Nathans. How to draw the line in biomedical research. eLife, 2, March 2013. ISSN 2050-084X. doi: 10.7554/eLife.00638. URL https://elifesciences.org/articles/00638.

Warrick J Inder, Goce Dimeski, and Anthony Russell. Measurement of salivary cortisol in 2012–laboratory techniques and clinical indications. Clinical endocrinology, 77(5):645–651, 2012.

Narayanan Kandasamy, Ben Hardy, Lionel Page, Markus Schaffner, Johann Graggaber, Andrew S Powlson, Paul C Fletcher, Mark Gurnell, and John Coates. Cortisol shifts financial risk preferences. Proc Natl Acad Sci U S A, 111(9):3608–3613, March 2014. doi: 10.1073/pnas.1317908111. URL http://www.pnas.org/cgi/doi/10.1073/pnas.1317908111.00000.

Jerker Karlén, Johnny Ludvigsson, Anneli Frostell, Elvar Theodorsson, and Tomas Faresjö. Cortisol in hair measured in young adults-a biomarker of major life stressors? BMC Clinical Pathology, 11(1):12, 2011.

KS Kendler, LM Karkowski, and CA Prescott. The assessment of dependence in the study of stressful life events: validation using a twin design. Psychological medicine, 29(06):1455–1460, 1999.

Lars Vedel Kessing, Esben Agerbo, and Preben B Mortensen. Does the impact of major stressful life events on the risk of developing depression change throughout life? Psychological medicine, 33(07):1177–1184, 2003.

Karolina M. Lempert, Anthony J. Porcelli, Mauricio R. Delgado, and Elizabeth Tricomi. Individual Differences in Delay Discounting Under Acute Stress: The Role of Trait Perceived Stress. Frontiers in Psychology, 3, July 2012. ISSN 1664-1078. doi: 10.3389/fpsyg.2012.00251. URL http://www.ncbi.nlm.nih.gov/pmc/articles/PMC3400439/.00037.

Karolina M. Lempert, Joseph T. McGuire, Danielle B. Hazeltine, Elizabeth A. Phelps, and Joseph W. Kable. The effects of acute stress on the calibration of persistence. Neurobiology of Stress, 8:1–9, February 2018a. ISSN 23522895. doi: 10.1016/j.ynstr.2017.11.001. URL https://linkinghub.elsevier.com/retrieve/pii/S2352289517300310.

Karolina M. Lempert, Joanna E. Steinglass, Anthony Pinto, Joseph W. Kable, and Helen Blair Simpson. Can delay discounting deliver on the promise of RDoC? Psychological Medicine, pages 1–10, August 2018b. ISSN 0033-2917, 1469-8978. doi: 10.1017/S0033291718001770. URL https://www.cambridge.org/core/product/identifier/S0033291718001770/type/journal_article.

Jennifer K. Lenow, Sara M. Constantino, Nathaniel D. Daw, and Elizabeth A. Phelps. Chronic and Acute Stress Promote Overexploitation in Serial Decision Making. The Journal of Neuroscience, 37(23):5681–5689, June 2017. ISSN 0270-6474, 1529-2401. doi: 10.1523/JNEUROSCI.3618-16.2017. URL http://www.jneurosci.org/lookup/doi/10.1523/JNEUROSCI.3618-16.2017.

Jennifer S Lerner, Ye Li, and Elke U Weber. The financial costs of sadness. Psychological science, page 0956797612450302, 2012.

Arnold Stanley Linsky and Murray Arnold Straus. Social stress in the United States: Links to regional patterns in crime and illness. Auburn House, 1986.

R. Linz, T. Singer, and V. Engert. Interactions of momentary thought content and subjective stress predict cortisol fluctuations in a daily life experience sampling study. Scientific Reports, 8(1):15462, December 2018. ISSN 2045-2322. doi: 10.1038/s41598-018-33708-0. URL http://www.nature.com/articles/s41598-018-33708-0.

Silvia Lopez-Guzman, Anna B. Konova, and Paul W. Glimcher. Computational psychiatry of impulsivity and risk: how risk and time preferences interact in health and disease. Philosophical Transactions of the Royal Society B: Biological Sciences, 374(1766):20180135, February 2019. ISSN 0962-8436, 1471-2970. doi: 10.1098/rstb.2018.0135. URL http://www.royalsocietypublishing.org/doi/10.1098/rstb.2018.0135.

Evgeniya Lukinova and Jeffrey C Erlich. Quantifying the contribution of individual variation in timing to delay-discounting. Scientific Reports, 11(1):1–13, 2021. doi: 10.1038/s41598-021-97496-w.

Evgeniya Lukinova, Yuyue Wang, Steven F Lehrer, and Jeffrey C Erlich. Time preferences are reliable across time-horizons and verbal versus experiential tasks. eLife, page 27, February 2019. doi: 10.7554/eLife.39656.001.

Gediminas Luksys and Carmen Sandi. Neural mechanisms and computations underlying stress effects on learning and memory. Current Opinion in Neurobiology, 21(3):502–508, June 2011. ISSN 09594388. doi: 10.1016/j.conb.2011.03.003. URL https://linkinghub.elsevier.com/retrieve/pii/S0959438811000432.

Scott M Lundberg and Su-In Lee. A unified approach to interpreting model predictions. Advances in neural information processing systems, 30, 2017.

Silvia U. Maier, Aidan B. Makwana, and Todd A. Hare. Acute Stress Impairs Self-Control in Goal-Directed Choice by Altering Multiple Functional Connections within the Brain’s Decision Circuits. Neuron, 87(3):621–631, August 2015. ISSN 08966273. doi: 10.1016/j.neuron.2015.07.005. URL https://linkinghub.elsevier.com/retrieve/pii/S0896627315006273.

James E Mazur. An adjusting procedure for studying delayed reinforcement. Commons, ML.; Mazur, JE.; Nevin, JA, pages 55–73, 1987.

Jennifer Meggs, J Golby, CJ Mallett, DF Gucciardi, and Remco CJ Polman. The Cortisol Awakening Response and Resilience in Elite Swimmers. International journal of sports medicine, 37(02):169–174, 2016.

Gregory E Miller, Edith Chen, and Eric S Zhou. If it goes up, must it come down? Chronic stress and the hypothalamic-pituitary-adrenocortical axis in humans. Psychological bulletin, 133(1):25, 2007.

Sendhil Mullainathan and Eldar Shafir. Scarcity: Why having too little means so much. Macmillan, 2013.

Fabio Paglieri. The costs of delay: Waiting versus postponing in intertemporal choice. Journal of the Experimental Analysis of Behavior, 99(3):362–377, 2013.

Jonathan W Peirce. PsychoPy—psychophysics software in Python. Journal of neuroscience methods, 162(1):8–13, 2007.

Vicente Prado-Gascó, Usue de la Barrera, Sandra Sancho-Castillo, José Enrique de la Rubia-Ortí, and Inmaculada Montoya-Castilla. Perceived stress and reference ranges of hair cortisol in healthy adolescents. PLOS ONE, 14(4):e0214856, April 2019. ISSN 1932-6203. doi: 10.1371/journal.pone.0214856. URL https://dx.plos.org/10.1371/journal.pone.0214856.

Marita Pruessner, Dirk H Hellhammer, Jens C Pruessner, and Sonia J Lupien. Self-reported depressive symptoms and stress levels in healthy young men: associations with the cortisol response to awakening. Psychosomatic Medicine, 65(1):92–99, 2003.

Michala Iben Riis-Vestergaard, Vanessa van Ast, Sandra Cornelisse, Marian Jöels, and Johannes Haushofer. The effect of hydrocortisone administration on intertemporal choice. Psychoneuroendocrinology, 88:173–182, February 2018. ISSN 03064530. doi: 10.1016/j.psyneuen.2017.10.002. URL http://linkinghub.elsevier.com/retrieve/pii/S0306453017305322.

Paul A Samuelson. A note on measurement of utility. The Review of Economic Studies, 4(2):155–161, 1937.

Robbie Schepers and C Rob Markus. Gene$ times$ cognition interaction on stress-induced eating: Effect of rumination. Psychoneuroendocrinology, 54:41–53, 2015.

Peter Sokol-Hessner, Candace M. Raio, Sarah P. Gottesman, Sandra F. Lackovic, and Elizabeth A. Phelps. Acute stress does not affect risky monetary decision-making. Neurobiology of Stress, 5:19–25, December 2016. ISSN 2352-2895. doi: 10.1016/j.ynstr.2016.10.003.00005.

Tobias Stalder, Susann Steudte-Schmiedgen, Nina Alexander, Tim Klucken, Annika Vater, Susann Wichmann, Clemens Kirschbaum, and Robert Miller. Stress-related and basic determinants of hair cortisol in humans: A meta-analysis. Psychoneuroendocrinology, 77:261–274, March 2017. ISSN 03064530. doi: 10.1016/j.psyneuen.2016.12.017. URL https://linkinghub.elsevier.com/retrieve/pii/S0306453016306114.

Taiki Takahashi. Cortisol levels and time-discounting of monetary gain in humans. Neuroreport, 15(13):2145–2147, September 2004. ISSN 0959-4965. 00063.

Berry J. van Holland, Monique H. W. Frings-Dresen, and Judith K. Sluiter. Measuring short-term and long-term physiological stress effects by cortisol reactivity in saliva and hair. International Archives of Occupational and Environmental Health, 85(8):849–852, November 2012. ISSN 0340-0131, 1432-1246. doi: 10.1007/s00420-011-0727-3. URL http://link.springer.com/10.1007/s00420-011-0727-3.

Barbara Vanaelst, Inge Huybrechts, Karin Bammann, Nathalie Michels, Tineke Vriendt, Krishna Vyncke, Isabelle Sioen, Licia Iacoviello, Kathrin Günther, Denes Molnar, Lauren Lissner, Noellie Rivet, Jean-Sebastien Raul, and Stefaan Henauw. Intercorrelations between serum, salivary, and hair cortisol and child-reported estimates of stress in elementary school girls: Cortisol intercorrelations and childhood stress. Psychophysiology, pages n/a–n/a, July 2012. ISSN 00485772. doi: 10.1111/j.1469-8986.2012.01396.x. URL http://doi.wiley.com/10.1111/j.1469-8986.2012.01396.x.

Carol Wang, Pelin Cunningham-Erdogdu, Mai-Ly Nguyen Steers, Andrew Phillip Weinstein, and Clayton Neighbors. Stressful life events and gambling: The roles of coping and impulsivity among college students. Addictive Behaviors, 107:106386, August 2020. ISSN 03064603. doi: 10.1016/j.addbeh.2020.106386. URL https://linkinghub.elsevier.com/retrieve/pii/S0306460319315497.

Lisa J. Weckesser, Friedericke Dietz, Kornelius Schmidt, Juliane Grass, Clemens Kirschbaum, and Robert Miller. The psychometric properties and temporal dynamics of subjective stress, retrospectively assessed by different informants and questionnaires, and hair cortisol concentrations. Scientific Reports, 9(1):1098, December 2019. ISSN 2045-2322. doi: 10.1038/s41598-018-37526-2. URL http://www.nature.com/articles/s41598-018-37526-2.

Quan Zhang, Zheng Chen, Shenghuo Chen, Tian Yu, Juxia Wang, Weiwen Wang, and Huihua Deng. Correlations of hair level with salivary level in cortisol and cortisone. Life Sciences, 193:57–63, January 2018. ISSN 00243205. doi: 10.1016/j.lfs.2017.11.037. URL https://linkinghub.elsevier.com/retrieve/pii/S0024320517306148.

