## Supplementary Information for "Willingness to wait covaries with endogenous variation in cortisol"

#### SI Results

##### General population group characteristics

The general population group was by large a homogeneous group representing adults from 22 to 49, with only a couple of subjects being older than 40 (Figure S1A). One might notice that among demographic variables the subject consumption level was spread a lot. This is due to some subjects counting the rent towards their consumption level as they reported in the end of the demographic survey. The body mass index (BMI) of male subjects was on average higher than of female subjects in the study, with most of the subjects having normal or healthy weight.

We used the Barratt Impulsiveness Scale (BIS-11; Patton et al., 1995) as a measure of impulsivity ($N = 40$). The mean total score for our sample was 63.35 (SD = 8.30, Figure S1B, similar to a student sample $M_{st} = 61.79$ in Lukinova et al. (2019) and previous literature). As before (Lukinova et al., 2019) we did not find any significant correlations between BIS-11 and discount factors. Among stress variables and deltas none were significantly correlated to BIS.

##### Fitting subjects' choices

The discount factors were estimated using a softmax-hyperbolic fit (i) in a similar way as in Lukinova et al. (2019), i.e. a four population level and three subject level parameters model - reduced model; (ii) with a noise per unit model; and (iii) with a reward scale model, where an additional parameter 'rews' scaled the delayed reward per unit. All mixed-effects models were done after conversion of all delays to days (resulted in the same fits as fits in the units of the task) with the 'brms' package in R (Bürkner, 2017).

The reduced model ($M_{4p,3s}$) had 4 population level parameters ($\log(k)$, and decision-noise, $\log(\tau)$ for each of the two delay-discounting tasks per session) and 3 parameters per subject: e.g., $\log(k_{SDD})$, $\log(k_{DDD})$ and $\log(\tau)$.

```
n_chose_later | trials(n_trials) ~
(later_reward/(1 + exp(logk)*delay)-sooner_reward)/exp(logtau),
logk ~ unit + (unit | subjid),
logtau ~ unit + (1 | subjid)
```

where `later_reward` is the later reward, `sooner_reward` is the sooner reward; `logk` is the natural logarithm of the discounting parameter $k$ and `logtau` ($\log(\tau)$) is the natural logarithm of the decision noise; in the binomial specification the data was grouped and summarized by distinct trial types, where `n_chose_later` is the count of choices when the later reward is selected and `n_trials` is the number of trials for a particular trial type.

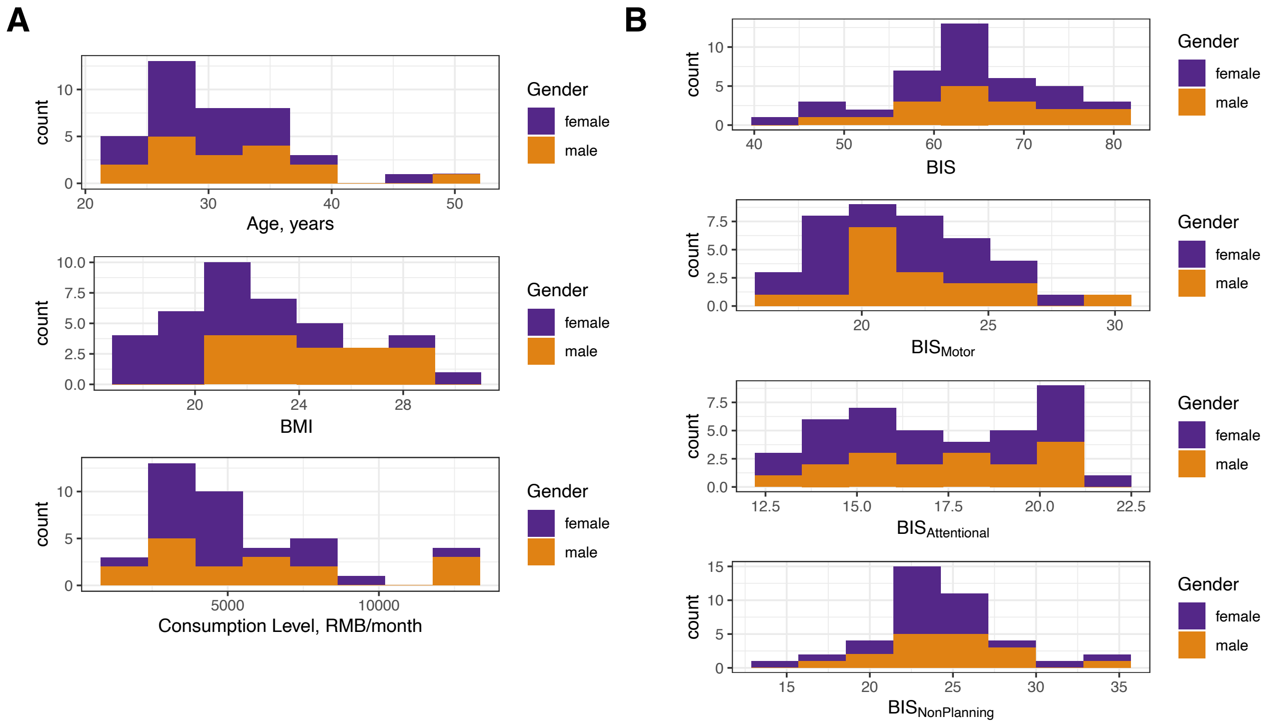


**Figure S1.** General Chinese population characteristics. (A) Stacked histograms of Age, BMI and consumption level (from top to bottom) divided by gender (color). (B) Stacked histograms of BIS and its components divided by gender (color). Note: the 'other' gender was masked to preserve anonymity.

The noise per unit model ($M_{4p,4s}$) had 4 population level parameters ($\log(k)$, and decision-noise, $\log(\tau)$ for each of the two delay-discounting tasks per session) and 4 parameters per subject: e.g., $\log(k_{SDD})$, $\log(k_{DDD})$, $\log(\tau_{SDD})$ and $\log(\tau_{DDD})$.

```
n_chose_later | trials(n_trials) ~
(later_reward/(1 + exp(logk)*delay)-sooner_reward)/exp(logtau),
logk ~ unit + (unit | subjid),
logtau ~ unit + (unit | subjid)
```

The main 'reward scale' model ($M_{6p,3s}$, model syntax provided in the main text) had 6 population level parameters ($\log(k)$, decision-noise, $\log(\tau)$, and reward scale parameter, rews, for each of the two delay-discounting tasks per session) and 3 parameters per subject: e.g., for session 1: $\log(k_{SDD})$, $\log(k_{DDD})$ and $\log(\tau)$. All model syntaxes were valid for fitting each session that includes two delay-discounting tasks each, where SDD (seconds delay discounting) and DDD (day delay discounting) should be changed to DDD and WDD (week delay discounting) for session 3.

The main model was better compared to the models that reflected the reduced number of parameters based on 10-fold cross validation criteria (Table S1). Therefore, 'reward scale' model (the main model) was used in the main text.

We reported the between tasks and between sessions Pearson correlations of subjects' discount factors ($\log(k)$) for all models under consideration in Table S2. The correlation coefficient between SDD and DDD tasks decreased with the increase in number of parameters in the model (nonsignificant difference according to 'cocor' tests done via R package 'cocor', (Diedenhofen and Musch, 2015)). The between-session correlation decreased slightly when reward scaling is used only between first and second sessions of the DDD task. Consistent with previous scholarship, we found high test/re-test reliability in delay discounting (with significant correlation coefficients for fits across sessions ranging from 0.52 to 0.87, reward scale model).

The subjects' choices in the decision-making sessions were well fit by the models. The Bayesian $r^2$ was reported in each of the subject plots (Figures S2 & S3 for sessions 1 & 2 combined with SDD and DDD, and

**Table S1.** Kfold model comparison

| | ΔELPD | ΔSE |
|---|---|---|
| **sessions 1 & 2** | | |
| reward scale | 0 | 0 |
| noise per unit | -439.8 | 60.2 |
| reduced | -483.8 | 56.5 |
| **session 1** | | |
| reward scale | 0 | 0 |
| noise per unit | -145.5 | 35.4 |
| reduced | -176.5 | 35.4 |
| **session 2** | | |
| reward scale | 0 | 0 |
| reduced | -277.5 | 40.5 |
| noise per unit | -288.6 | 48.3 |
| **session 3** | | |
| reward scale | 0 | 0 |
| noise per unit | -18.5 | 21.4 |
| reduced | -27.5 | 18.6 |

*Note.* 10-fold cross validation was done using the `kfold` function in the `brms` R package. The `loo_compare` function in the `brms` package arranges the models in order, from best to worst. The second column showed the difference between the expected log pointwise predictive density (ΔELPD, Vehtari et al., 2017) of the best model with the model on that row.

for session 3 with DDD and WDD, respectively). The posterior plots for the fits were shown in Figure S4.

##### Validating Stress Measures

The summary statistics of stress measures corresponded well to the expected values from the literature. All means and standard deviations are reported in the Table S3. As such, our participants PSS scores corresponded to low and moderate stress levels (0-26), while BEPSI five close-ended questions (0-4 for low and moderate stress per question compared to our mean of $25/5 = 5$ per question) and LCUs (150-299 $\sim$ a 50% chance of suffering from stress) indicated several participants to be at a higher stress level . As for cortisol levels, saliva measures were well below reference values in the afternoon for disease screening - 4 ng/mL. Similarly, hair cortisol levels for most of our participants were well below the reference median of 46.1 pg/mg (Sauvé et al., 2007).

Many scholars practice log-transformation of stress variables due to skewed distributions of the raw values, among them Lenow et al. (2017) and Lempert et al. (2018). Our data also showed that pattern in Figure S5A. That is why, in the main text we presented results after such transformation (Figure S5B). Although some researchers also control for gender (Takahashi, 2004; Barel et al., 2017), our log-transformed stress data as well as questionnaires did not show any main effect of gender (e.g., stress deltas after log-transformation split by gender in Figure S5C-D).

We also conducted the principal component analysis (PCA) to visualize the relationships across the stress modalities. To this end, we used the nine stress-vectors (three saliva, three hair, and three questionnaire data points) for each subject in Figure S6. Above 70% of variance was explained by the first four principle components. The respective largest loadings per principle component were shown in Figure S6C inset. The second principle component along the nine stress-vectors distinguished between human bio samples (hair and saliva) and questionnaires, whereas the third principle component distinguished between hair and saliva (Figure S6B). According to the angles (calculated from the positive x-axis following the four quadrants of 2D plane) in Table S4 we got additional evidence that clusters of stress modalities were different. We used the first four principal components from this analysis for correlation analysis.

Some researchers (Lenow et al., 2017; Lempert et al., 2018) also practice outlier removal from cortisol data. Following their procedure, two participants should be removed from analyses due to an abnormally high

**Table S2.** Correlations of subjects' discount factors

| | Reduced Model | Noise Per Unit Model | Reward Scale Model | N |
|---|---|---|---|---|
| **between tasks** | | | | |
| session 1 (SDD1 vs. DDD1) | $r = 0.52$ | $r = 0.53$ | $r = 0.50$ | 41 |
| | $p = 0.0005$ | $p = 0.0004$ | $p = 0.0008$ | |
| session 2 (SDD2 vs. DDD2) | $r = 0.47$ | $r = 0.50$ | $r = 0.48$ | 40 |
| | $p = 0.0022$ | $p = 0.0010$ | $p = 0.0019$ | |
| session 3 (WDD vs. DDD3) | $r = 0.97$ | $r = 0.97$ | $r = 0.96$ | 41 |
| | $p < 2.2e^{-16}$ | $p < 2.2e^{-16}$ | $p < 2.2e^{-16}$ | |
| **between sessions** | | | | |
| SDD1 vs. SDD2 | $r = 0.60$ | $r = 0.60$ | $r = 0.66$ | 40 |
| | $p = 4.684e^{-05}$ | $p = 4.882e^{-05}$ | $p = 3.96e^{-06}$ | |
| DDD1 vs. DDD2 | $r = 0.53$ | $r = 0.56$ | $r = 0.52$ | 40 |
| | $p = 0.0004$ | $p = 0.0002$ | $p = 0.0006$ | |
| DDD2 vs. DDD3 | $r = 0.89$ | $r = 0.88$ | $r = 0.87$ | 40 |
| | $p = 2.374e^{-14}$ | $p = 1.526e^{-13}$ | $p = 2.974e^{-13}$ | |
| DDD1 vs. DDD3 | $r = 0.54$ | $r = 0.54$ | $r = 0.54$ | 41 |
| | $p = 0.0002$ | $p = 0.0002$ | $p = 0.0003$ | |

*Note.* In session 2, one subject was not sensitive to delay and was excluded from that session fit (reflected in N).

**Table S3.** Stress Measures Summary Statistics

| | Mean | Standard Deviation |
|---|---|---|
| salivary cortisol, ng/mL | 0.925 | 0.608 |
| hair cortisol, pg/mg | 17.839 | 12.475 |
| PSS | 16.368 | 6.105 |
| BEPSI | 25.105 | 9.426 |
| LCU | 281.947 | 161.296 |

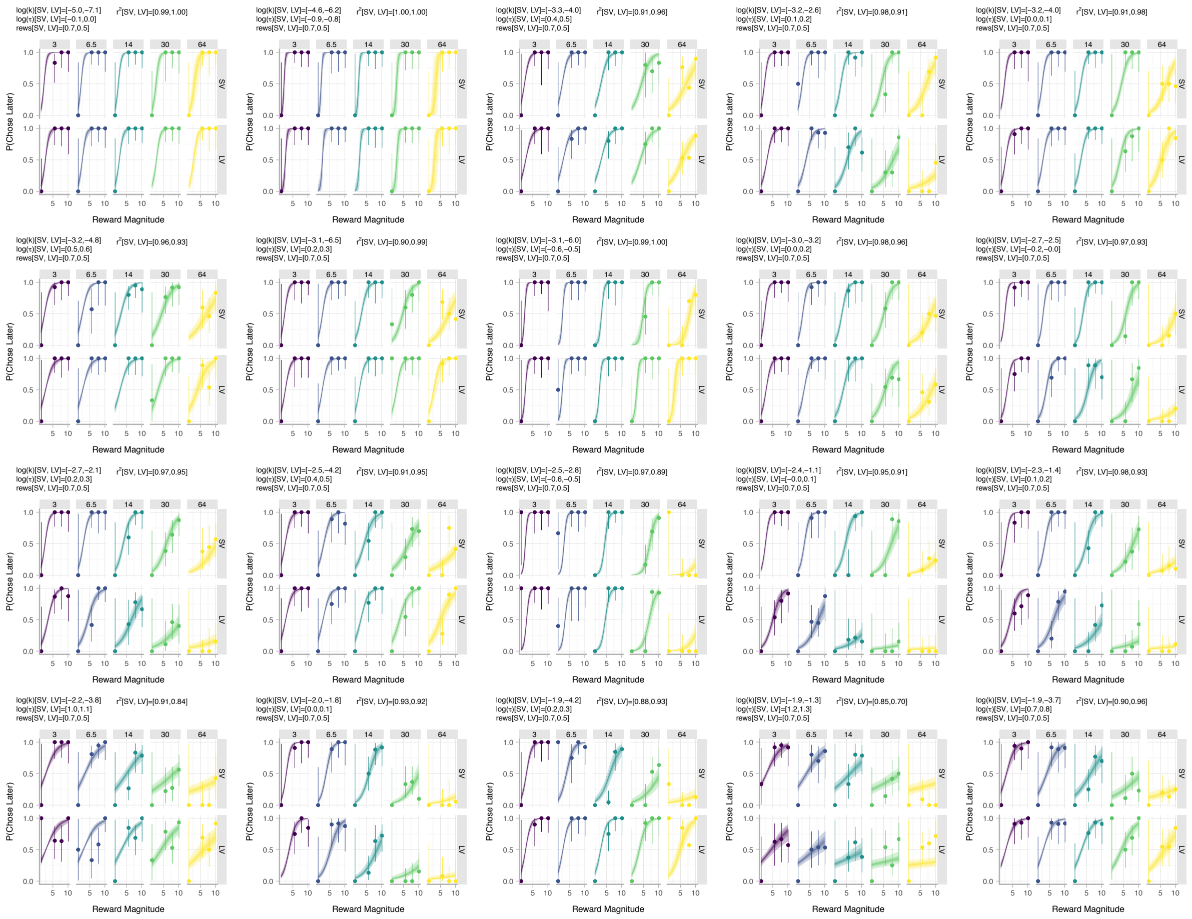

**Figure S2.** In this figure SV is SDD and LV is DDD. Subject plots were arranged by $\log(k_{SV})$ (from smaller to larger). In each panel, the marker and error bar indicate the mean and binomial confidence intervals of the subject's choices for that offer. The facets (columns) and colors signify the delay of the later reward in the units of the task. The smooth ribbon indicated the BHM model fits (at 50, 80, 99% credible intervals). At the top of the subject plot we indicate the mean estimates of $\log(k)$, $\log(\tau)$ and rews for each task for that subject. We also indicate the Bayesian $r^2$ for each task. Multi-page PDF, available as a file at https://github.com/erlichlab/gpstress.

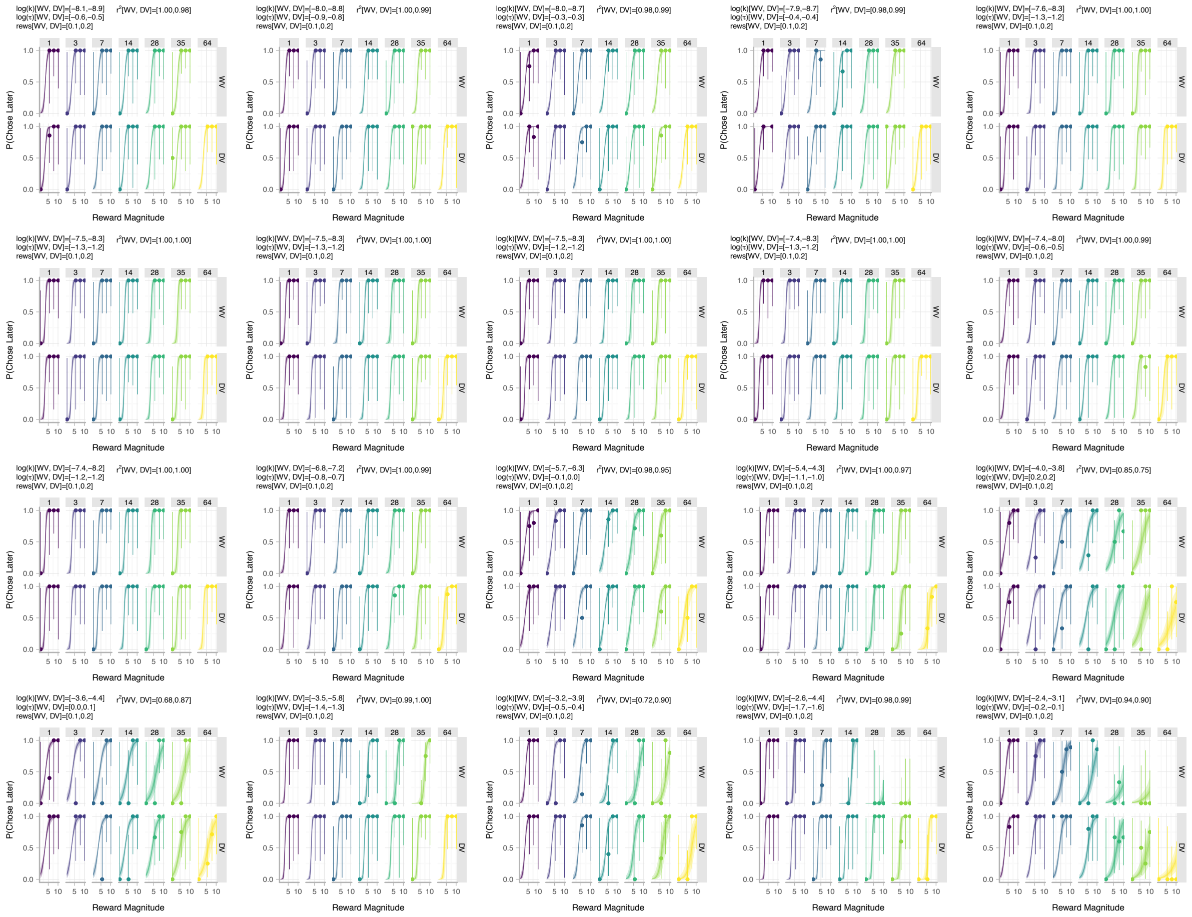

**Figure S3.** In this figure DV is DDD and WV is WDD. Subject plots were arranged by $\log(k_{WV})$ (from smaller to larger). In each panel, the marker and error bar indicate the mean and binomial confidence intervals of the subject's choices for that offer. The facets (columns) and colors signify the delay of the later reward in the units of the task. The smooth ribbon indicated the BHM model fits (at 50, 80, 99% credible intervals). At the top of the subject plot we indicate the mean estimates of $\log(k)$, $\log(\tau)$ and rews for each task for that subject. We also indicate the Bayesian $r^2$ for each task. Multi-page PDF, available as a file at https://github.com/erlichlab/gpstress.

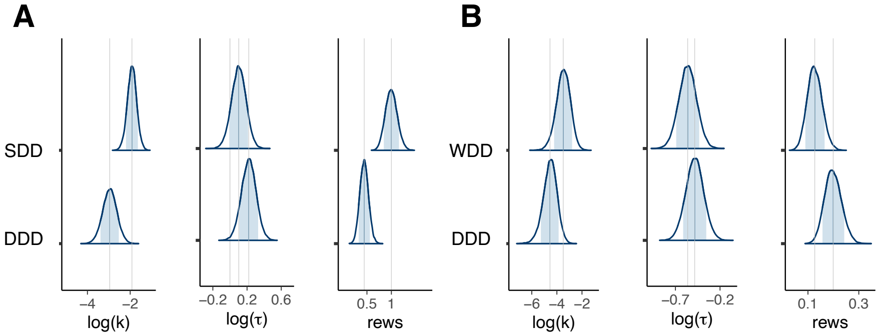


**Figure S4.** Distributions of posterior parameter estimates of population-level parameters from the model fits: (A) discount factor, $\log(k)$, decision noise, $\log(\tau)$, and reward scale parameter, rews, for the SDD and DDD tasks; (B) $\log(k)$, $\log(\tau)$ and rews for the WDD and DDD tasks.

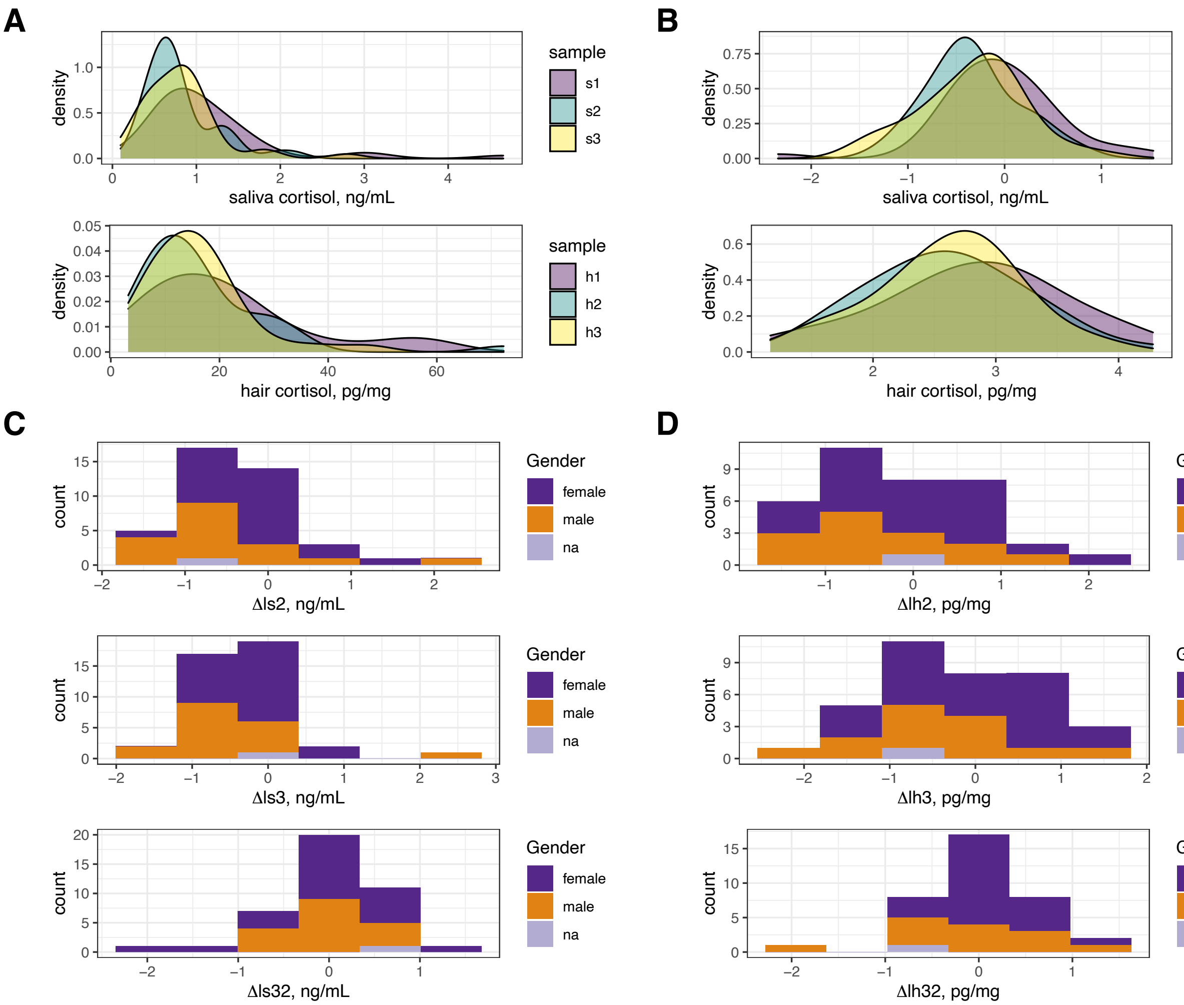


**Figure S5.** (A-B) Distributions of stress variables before (A) and after log-transformation (B): first row - salivary cortisol and second row - hair cortisol. Each facet has a kernel density estimate of a stress variable identified by color label per each facet. (C-D) Stacked histograms of cortisol deltas after log-transformation divided by gender (color). (C) Salivary cortisol delta: Δls. (D) Hair cortisol delta: Δlh.

A

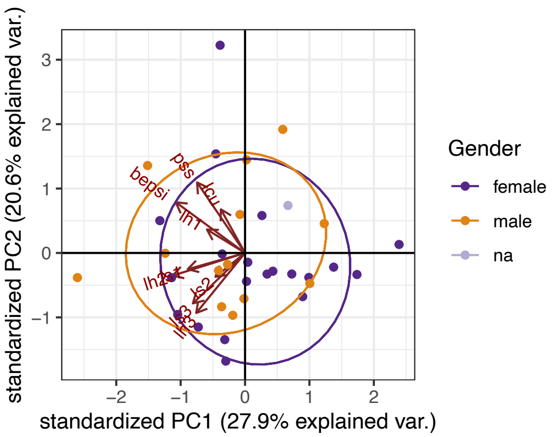


B

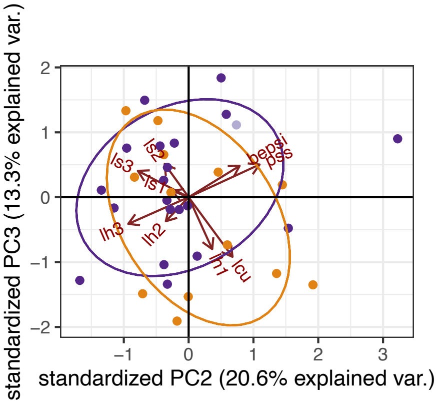


C

| PC # | Three largest loadings | | |
|---|---|---|---|
| PC1 | lh2 | bepsi | ls1 |
| PC2 | pss | lh3 | bepsi |
| PC3 | lcu | lh1 | ls2 |
| PC4 | ls2 | lh1 | ls3 |

**Figure S6.** N = 34 (A-B) Two 2D planes with axes being three first principle components (PCs): (A) PC1-PC2 plane, and (B) PC2-PC3 plane. These are PCA biplots that combine standardized data and the stress variables loadings (the arrows, with directions of corresponding eigenvectors). The data color is based on the gender group. The ellipses surrounding the data points (with color corresponding to the gender group) are drawn around clusters identified after automatic classification, based on PCA scores. (C) A table of the respective largest three loadings per principle component, in order. Large (either positive or negative) loadings indicate that this stress variable has a strong effect on that principal component.

**Table S4.** PCA angles

| | PC1-PC2 plane | PC2-PC3 plane |
|---|---|---|
| ls1 | 196° | 159° |
| ls2 | 219° | 126° |
| ls3 | 224° | 153° |
| lh1 | 148° | 295° |
| lh2 | 197° | 227° |
| lh3 | 231° | 204° |
| pss | 124° | 25° |
| bepsi | 144° | 32° |
| lcu | 119° | 306° |

baseline cortisol concentration of saliva (log(s1)) that was > 2.3 standard deviations from the mean (Figure S7). Since we did not have many significant results associated with saliva sample cortisol level, we decided not to re-run our analysis without these outliers.

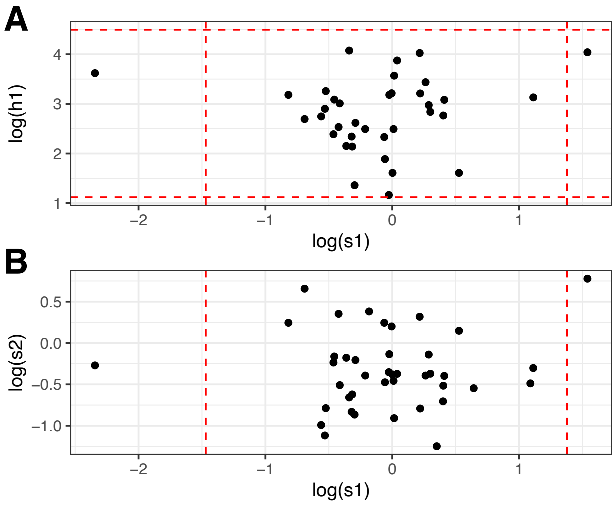


**Figure S7.** (A-B) Baseline stress outlier check. Red-dashed lines signify the 2.3 standard deviations away from the mean. Each dot is one subject. (A) log(s1) (x-axis) against log(h1) (y-axis, $N = 36$); (B) log(s1) (x-axis) against log(s2) (y-axis, $N = 41$).

#### Additional stress/behavior analysis

We distinguished between within-subject and between-subject analysis. Therefore, we created two tables that contain all correlations performed (within-subject in Table S5 and between-subject in Table S6).

We considered not only model-based delay-discounting parameters for the correlation analysis, but also model-free measures. These include subjects' profit, total waiting time (wait), proportion of later choices (PL), reaction time (RT), and their change. It is important to note that the total waiting time and the profit variables were experienced only in the seconds delay task (SDD) and in the other tasks only represent potential outcomes (hypothetical total) compared to only one random outcome selected for payout. Again for the model-free measures we find correlations only with more 'real' seconds task. For example, changes in hair cortisol were strongly and negatively associated with changes in choosing later option in SDD ($\Delta PL_{S_2} = PL_{S_2} - PL_{S_1}$ vs. Δlh2: Pearson $r = -0.34$, $p = 0.046$; $\Delta PL_{S_2}$ vs. Δlh3: Pearson $r = -0.49$, $p = 0.003$, with the latter staying significant after adjusting for multiple comparisons) further supporting results in the main text.

If endogenous cortisol has been shown to reliably reflect the activity of the hypothalamic-pituitary-adrenal

**Table S5.** Within-subject correlation analysis

| | $\log(k_{SDD2}) - \log(k_{SDD1})$ | $PL_{S_2} - PL_{S_1}$ | $\log(k_{DDD2}) - \log(k_{DDD1})$ | $\log(k_{DDD3}) - \log(k_{DDD1})$ | $\log(k_{DDD3}) - \log(k_{DDD2})$ |
|---|---|---|---|---|---|
| **Salivary cortisol:** $N = 41$ | | | | | |
| $\Delta$ls2 = | $r = 0.16$ | $r = -0.01$ | $r = -0.03$ | $r = 0.10$ | $r = 0.22$ |
| = log(s2)−log(s1) | $p = 0.321$ | $p = 0.974$ | $p = 0.863$ | $p = 0.524$ | $p = 0.170$ |
| $\Delta$ls3 = | $r = -0.09$ | $r = 0.13$ | $r = -0.19$ | $r = -0.07$ | $r = 0.15$ |
| = log(s3)−log(s1) | $p = 0.601$ | $p = 0.413$ | $p = 0.244$ | $p = 0.687$ | $p = 0.350$ |
| $\Delta$ls32 = | $r = -0.27$ | $r = 0.15$ | $r = -0.17$ | $r = -0.19$ | $r = -0.09$ |
| = log(s3)−log(s2) | $p = 0.087$ | $p = 0.357$ | $p = 0.291$ | $p = 0.245$ | $p = 0.592$ |
| **Hair cortisol:** $N = 36$ | | | | | |
| $\Delta$lh2 = | $r = 0.17$ | $r =$ **-0.34** | $r = 0.09$ | $r = 0.00$ | $r = -0.14$ |
| = log(h2)−log(h1) | $p = 0.318$ | $p =$ **0.046** | $p = 0.596$ | $p = 0.979$ | $p = 0.428$ |
| $\Delta$lh3 = | $r =$ **0.36** | $r =$ **-0.49** | $r = 0.23$ | $r = 0.05$ | $r = -0.24$ |
| = log(h3)−log(h1) | $p =$ **0.033** | $p =$ **0.003** | $p = 0.175$ | $p = 0.793$ | $p = 0.151$ |
| $\Delta$lh32 = | $r = 0.30$ | $r = -0.27$ | $r = 0.22$ | $r = 0.07$ | $r = -0.18$ |
| = log(h3)−log(h2) | $p = 0.079$ | $p = 0.118$ | $p = 0.198$ | $p = 0.669$ | $p = 0.302$ |

*Note.* Significant correlations at .05 level were highlighted.

**Table S6.** Between-subject correlation analysis

| | $\log(k_{SDD1})$ | $PL_{S_1}$ | $\log(k_{SDD2})$ | $PL_{S_2}$ | $\log(k_{DDD1})$ | $\log(k_{DDD2})$ | $\log(k_{DDD3})$ |
|---|---|---|---|---|---|---|---|
| **Salivary cortisol:** $N = 41$ | | | | | | | |
| log(s1) | $r = -0.05$ | $r = 0.18$ | $r = -0.01$ | $r = 0.06$ | $r = 0.05$ | $r = 0.12$ | $r = -0.06$ |
| | $p = 0.738$ | $p = 0.273$ | $p = 0.948$ | $p = 0.720$ | $p = 0.774$ | $p = 0.473$ | $p = 0.692$ |
| log(s2) | $r = -0.29$ | $r = 0.18$ | $r = -0.03$ | $r = 0.02$ | $r = -0.03$ | $r = 0.03$ | $r = 0.10$ |
| | $p = 0.075$ | $p = 0.266$ | $p = 0.837$ | $p = 0.922$ | $p = 0.843$ | $p = 0.845$ | $p = 0.542$ |
| log(s3) | $r = 0.04$ | $r = 0.03$ | $r = 0.00$ | $r = 0.03$ | $r = 0.25$ | $r = 0.08$ | $r = 0.03$ |
| | $p = 0.813$ | $p = 0.850$ | $p = 0.990$ | $p = 0.843$ | $p = 0.113$ | $p = 0.633$ | $p = 0.867$ |
| **Hair cortisol:** $N = 36$ | | | | | | | |
| log(h1) | $r = 0.28$ | $r = -0.33$ | $r = 0.10$ | $r = -0.06$ | $r = 0.01$ | $r = -0.11$ | $r = -0.03$ |
| | $p = 0.099$ | $p = 0.053$ | $p = 0.557$ | $p = 0.711$ | $p = 0.962$ | $p = 0.538$ | $p = 0.881$ |
| log(h2) | $r = 0.24$ | $r = -0.17$ | $r = 0.23$ | $r = -0.25$ | $r = 0.21$ | $r = 0.17$ | $r = 0.10$ |
| | $p = 0.160$ | $p = 0.326$ | $p = 0.177$ | $p = 0.141$ | $p = 0.209$ | $p = 0.334$ | $p = 0.577$ |
| log(h3) | $r = -0.24$ | $r = 0.32$ | $r = -0.02$ | $r = -0.03$ | $r = -0.02$ | $r = 0.19$ | $r = 0.01$ |
| | $p = 0.152$ | $p = 0.058$ | $p = 0.920$ | $p = 0.868$ | $p = 0.922$ | $p = 0.265$ | $p = 0.932$ |
| **Questionnaires:** $N = 38$ | | | | | | | |
| PSS | $r = 0.30$ | $r = -0.23$ | $r = 0.16$ | $r = -0.08$ | $r = 0.01$ | $r = -0.04$ | $r = 0.02$ |
| | $p = 0.069$ | $p = 0.172$ | $p = 0.359$ | $p = 0.616$ | $p = 0.943$ | $p = 0.827$ | $p = 0.920$ |
| BEPSI | $r = 0.25$ | $r = -0.24$ | $r = 0.13$ | $r = -0.13$ | $r = 0.02$ | $r = 0.05$ | $r = 0.05$ |
| | $p = 0.139$ | $p = 0.138$ | $p = 0.431$ | $p = 0.452$ | $p = 0.906$ | $p = 0.770$ | $p = 0.759$ |
| LCU | $r = 0.24$ | $r = -0.31$ | $r = 0.29$ | $r = -0.31$ | $r = 0.08$ | $r = 0.01$ | $r = 0.10$ |
| | $p = 0.149$ | $p = 0.056$ | $p = 0.085$ | $p = 0.062$ | $p = 0.653$ | $p = 0.947$ | $p = 0.549$ |
| **PCs:** $N = 34$ | | | | | | | |
| PC1 | $r = 0.05$ | $r = -0.15$ | $r = -0.01$ | $r = 0.07$ | $r = -0.19$ | $r = -0.20$ | $r = -0.01$ |
| | $p = 0.784$ | $p = 0.382$ | $p = 0.955$ | $p = 0.683$ | $p = 0.276$ | $p = 0.268$ | $p = 0.972$ |
| PC2 | $r =$ **0.52** | $r =$ **-0.52** | $r = 0.31$ | $r = -0.30$ | $r = 0.15$ | $r = 0.08$ | $r = 0.14$ |
| | $p =$ **0.002** | $p =$ **0.002** | $p = 0.074$ | $p = 0.088$ | $p = 0.398$ | $p = 0.669$ | $p = 0.445$ |
| PC3 | $r = 0.32$ | $r =$ **-0.34** | $r = 0.11$ | $r = -0.06$ | $r = 0.14$ | $r = -0.09$ | $r = 0.06$ |
| | $p = 0.069$ | $p =$ **0.050** | $p = 0.540$ | $p = 0.749$ | $p = 0.429$ | $p = 0.599$ | $p = 0.757$ |
| PC4 | $r = 0.22$ | $r = -0.22$ | $r = 0.00$ | $r = -0.02$ | $r = 0.15$ | $r = -0.16$ | $r = -0.17$ |
| | $p = 0.211$ | $p = 0.218$ | $p = 0.991$ | $p = 0.918$ | $p = 0.407$ | $p = 0.376$ | $p = 0.345$ |

*Note.* Significant correlations at .05 level were highlighted.

(HPA) axis, the ratio of cortisol to cortisone (Rcc) is the potential biomarker to reliably assess the activity of 11$\beta$-hydroxysteriod dehydrogenases (11$\beta$-HSD) (Zhang et al., 2017). Cortisol levels together with cortisone levels for hair samples were available as a result of the LC-MS/MS atmospheric pressure chemical ionization analysis performed by Dr. Huihua Deng's lab at Southeast University (Chen et al., 2013). The ratio of cortisol to cortisone (Rcc) was not correlated with any of the model-based behavioral measures. However, we found two significant relationships of cortisol-to-cortisone ratio with model-free measures that lose significance after Bonferroni multiple comparison correction. Rcc1 (baseline ratio value that corresponds to two months before collection) was negatively correlated with profit in session 1 of the seconds delay task (Pearson $r = -0.36$, $p = 0.034$) and logarithmic transformation of Rcc2 (ratio value that corresponds to one month before collection) was negatively correlated with proportion of later choices in session 1 of the day delay task (Pearson $r = -0.37$, $p = 0.027$).

Following Lempert et al. (2018) we checked whether we can find any curvilinear relationships for those with increase in biological stress level compared to the baseline. There were only 13 subjects with such positive changes from lh1 to lh2 ($\Delta$lh2), 16 subjects from lh1 to lh3 ($\Delta$lh3) and 17 subjects from lh2 to lh3 ($\Delta$lh32). Out of many possible comparisons we found just a few inverted-U-shaped relationships between hair stress deltas and model-free parameters. All visually salient curvilinear relationships were plotted in Figure S8. Only one of them was close to be preferred using likelihood ratio tests (by checking the change in likelihood using a $\chi^2$ statistic). The complex model (polynomial of degree 2) could be preferred to the nested one at .1 level of significance (behavior $\sim \Delta$stress) for:

RT_S $\sim \Delta lh2 + (\Delta lh2)^2$, $P(> \chi^2) = 0.054$, $\beta((\Delta lh2)^2) = -0.62$, $p = 0.100$.

### References


Efrat Barel, Shosh Shahrabani, and Orna Tzischinsky. Sex Hormone/Cortisol Ratios Differentially Modulate Risk-Taking in Men and Women. *Evolutionary Psychology*, 15(1):147470491769733, January 2017. ISSN 1474-7049, 1474-7049. doi: 10.1177/1474704917697333. URL http://journals.sagepub.com/doi/10.1177/1474704917697333.

Paul-Christian Bürkner. brms: An R Package for Bayesian Multilevel Models Using Stan. *Journal of Statistical Software*, 80(1):1–28, 2017. doi: 10.18637/jss.v080.i01.

Zheng Chen, Jifeng Li, Jing Zhang, Xue Xing, Wei Gao, Zuhong Lu, and Huihua Deng. Simultaneous determination of hair cortisol, cortisone and DHEAS with liquid chromatography–electrospray ionization-tandem mass spectrometry in negative mode. *Journal of Chromatography B*, 929:187–194, June 2013. ISSN 15700232. doi: 10.1016/j.jchromb.2013.04.026. URL https://linkinghub.elsevier.com/retrieve/pii/S1570023213002365.

Birk Diedenhofen and Jochen Musch. cocor: A Comprehensive Solution for the Statistical Comparison of Correlations. *PLOS ONE*, 10(4):e0121945, April 2015. ISSN 1932-6203. doi: 10.1371/journal.pone.0121945. URL http://dx.plos.org/10.1371/journal.pone.0121945.

Karolina M. Lempert, Joseph T. McGuire, Danielle B. Hazeltine, Elizabeth A. Phelps, and Joseph W. Kable. The effects of acute stress on the calibration of persistence. *Neurobiology of Stress*, 8:1–9, February 2018. ISSN 23522895. doi: 10.1016/j.ynstr.2017.11.001. URL https://linkinghub.elsevier.com/retrieve/pii/S2352289517300310.

Jennifer K. Lenow, Sara M. Constantino, Nathaniel D. Daw, and Elizabeth A. Phelps. Chronic and Acute Stress Promote Overexploitation in Serial Decision Making. *The Journal of Neuroscience*, 37(23):5681–5689, June 2017. ISSN 0270-6474, 1529-2401. doi: 10.1523/JNEUROSCI.3618-16.2017. URL http://www.jneurosci.org/lookup/doi/10.1523/JNEUROSCI.3618-16.2017.

Evgeniya Lukinova, Yuyue Wang, Steven F Lehrer, and Jeffrey C Erlich. Time preferences are reliable across time- horizons and verbal versus experiential tasks. *eLife*, page 27, February 2019. doi: https://doi.org/10.7554/eLife.39656.001.

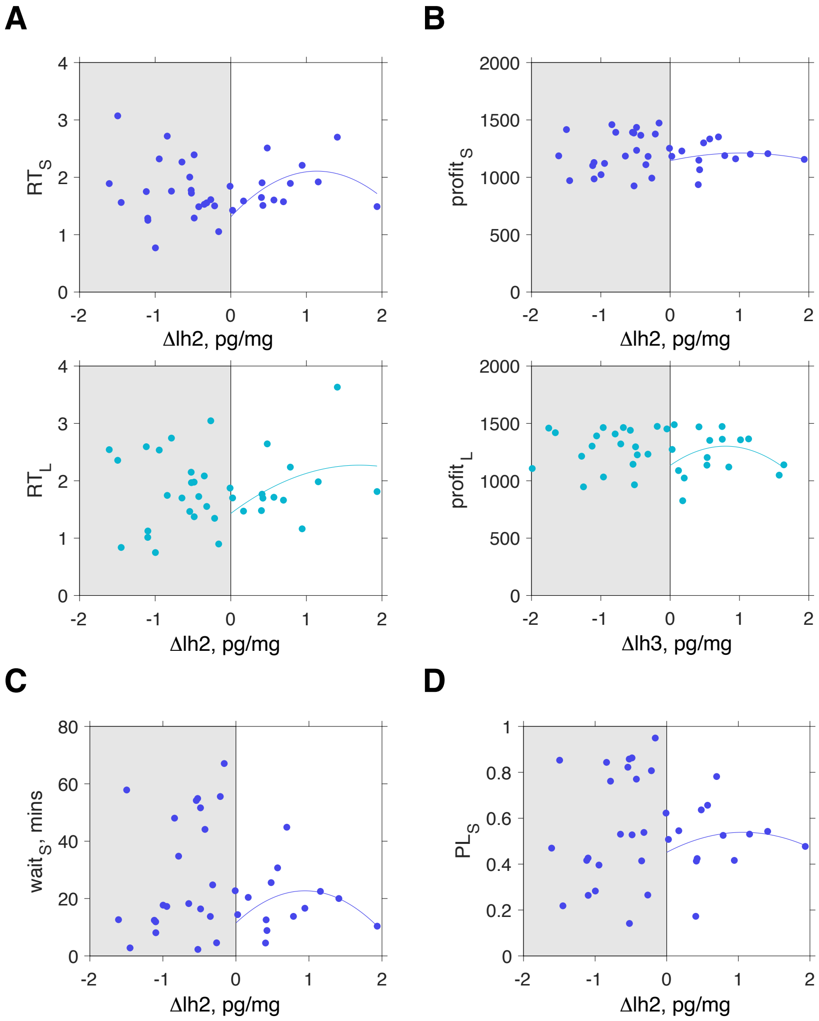


**Figure S8.** Curvilinear relationship between positive stress deltas (after log-transformation), model-free and other task measures. The polynomial of degree 2 was fit only to the stress deltas $> 0$. (A) Reaction time in seconds (S) and day (L) delay tasks. (B) Profit in seconds (S) and day (L) delay tasks. (C) Total wait time in SDD. (D) Proportion of later choices in SDD.

Jim H Patton, Matthew S Stanford, and others. Factor structure of the Barratt impulsiveness scale. *Journal of clinical psychology*, 51(6):768–774, 1995.

Brittany Sauvé, Gideon Koren, Grace Walsh, Sonya Tokmakejian, and Stan HM Van Uum. Measurement of cortisol in human hair as a biomarker of systemic exposure. *Clinical and Investigative Medicine*, pages E183–E191, 2007.

Taiki Takahashi. Cortisol levels and time-discounting of monetary gain in humans. *Neuroreport*, 15(13): 2145–2147, September 2004. ISSN 0959-4965. 00063.

Aki Vehtari, Andrew Gelman, and Jonah Gabry. Practical Bayesian model evaluation using leave-one-out cross-validation and WAIC. *Statistics and Computing*, 27(5):1413–1432, September 2017. ISSN 0960-3174, 1573-1375. doi: 10.1007/s11222-016-9696-4. URL `https://link.springer.com/article/10.1007/s11222-016-9696-4`. 00084.

Quan Zhang, Zheng Chen, Shenghuo Chen, Youyun Xu, and Huihua Deng. Intraindividual stability of cortisol and cortisone and the ratio of cortisol to cortisone in saliva, urine and hair. *Steroids*, 118:61–67, February 2017. ISSN 0039128X. doi: 10.1016/j.steroids.2016.12.008. URL `https://linkinghub.elsevier.com/retrieve/pii/S0039128X16301969`.
